# Flexible, task-dependent bimanual coordination along movement direction and extent

**DOI:** 10.1101/2025.09.24.678336

**Authors:** Preethi Ravikumar, Pratik K. Mutha

## Abstract

Recent work suggests that coordination of bilateral arm movements is mediated through flexible, task-dependent control policies rather than coupled motor commands that activate homologous muscle groups in the two arms. Here we examined whether such flexibility in bimanual control extends independently across movement direction and extent. We employed a bimanual reach task in which we altered the contribution of each arm to the perpendicular (direction axis) and/or parallel (extent axis) position of a single feedback cursor controlled by both arms together. We first replicated findings of Kitchen et al. (2023) showing that when one arm contributed more to perpendicular cursor motion, its lateral variability was reduced, while variability of the other arm increased. We then extended this result to perturbations affecting movement extent, observing similar asymmetric adjustments in variability in the parallel direction. Subsequent manipulations where contributions along both axes were manipulated simultaneously, revealed that irrespective of the gain combinations, the arm with the higher perpendicular contribution showed reduced lateral variability and the arm with the higher parallel contribution demonstrated restricted parallel variability, while allowing the corresponding lower contribution arms to compensate for perturbation-induced errors. These results suggest that: 1) the sensorimotor system always prioritizes corrections for deviations directly impacting task goals while tolerating higher variability in less relevant dimensions, and 2) that it is capable of adjusting coordination along movement direction and extent largely independently based on peripheral task demands. These findings add to the growing body of evidence supporting task-dependent modulation of bimanual motor coordination.

**SIGNIFICANCE STATEMENT:** Bimanual actions in humans are thought to be controlled via a control policy that can be flexibly tuned to task demands. This work aimed to demonstrate that such flexibility in control extends independently to movement direction and extent. We used a shared-cursor bimanual reaching task to selectively manipulate the contribution of each arm to cursor motion along the direction and extent axes first in isolation, then simultaneously. Our results show that the sensorimotor system restricts variability in the more task-relevant arm along each axis, enabling axis-specific coordination. These findings thus provide evidence that bimanual coordination along movement direction and extent can be governed flexibly and largely independently in response to changing task demands.

## INTRODUCTION

The human motor system exhibits remarkable adaptability in coordinating arm movements across a wide variety of tasks, ranging from applauding to holding a cup while texting. This versatility illustrates the ability of the sensorimotor system to organize and execute both independent as well as coordinated movements of the limbs. A large body of research has investigated the mechanisms and computations underlying coordinated or concurrent movements of the two upper limbs, i.e. bimanual coordination. This work has generally revealed that when individuals attempt to simultaneously perform different tasks with the two arms, such as drawing a circle with one hand and a line with the other (Franz et al. 1991; Franz, 1997), the movements of the arms end up being more symmetric even though the task demands asymmetry (Kelso et al., 1979b; Kelso et al, 1983; Marteniuk et al., 1984). This tendency to switch to more spatiotemporally symmetric movements has led to the view that upper limb bimanual coordination is primarily mediated by coupled motor commands that constrain homologous muscle groups across the two arms to “act as a single unit” (Kelso et al., 1979a).

If the above coupling hypothesis is true, then perturbations or task changes applied to one limb should elicit identical responses in both limbs. However, this idea has not been well supported in the literature (Diedrichsen, 2007; Diedrichsen and Gush, 2009; Mutha and Sainburg, 2009; Dimitriou et al., 2012; Omrani et al., 2013; Ranganathan et al., 2019; Kitchen et al. 2023; Yuk et al., 2024). Broadly, studies have shown that the responses observed in the two arms are contingent upon task conditions and goals rather than spatiotemporal symmetry alone. For example, both Diedrichsen (2007) and Mutha and Sainburg (2009) showed in the context of bimanual motion, that when each arm independently controlled the motion of a feedback cursor towards its own target, perturbations applied to one arm did not elicit error corrections in the unperturbed arm. Conversely, when both arms drove a single, shared cursor to a common target, perturbations applied to one arm did in fact result in corrections in the unperturbed arm. This differential presence of cross-limb interactions suggests a more flexible control strategy during bimanual reaching that is tied closely to the task goal, rather than a rigid “coupled” control process.

Recently, Kitchen et al. (2023) took this idea a step further by demonstrating that even when both arms are required to collaborate to achieve the task goal (by driving a single shared cursor to a single target), there remains remarkable flexibility in how each arm responds to task contingencies. These authors differentially varied the contribution of each arm to the horizontal motion of a shared feedback cursor during bilateral forward reaching. Specifically, a differential perpendicular gain was imposed on the cursor such that small deviations away from the ideal straight-line motion of each arm would result in amplified errors for one arm and attenuated errors for the other. This “perturbation” imposed strict positional constraints on the arms since any uncompensated deviations perpendicular to the movement trajectory would result in the cursor missing the target. Kitchen et al. (2023) observed that participants countered the perturbation by curtailing motor variability in the arm with the higher contribution to the perpendicular motion of the cursor, while allowing the other arm (that contributed less to the cursor’s perpendicular motion) to vary more freely. This asymmetric adjustment of lateral variability pointed to a flexible, task-dependent modulation within each arm even though both arms were collaborating to achieve a shared goal.

Intriguingly, Kitchen and colleagues varied only the perpendicular gain on the cursor while leaving the parallel contributions (contribution of each arm to cursor motion in the direction of target) matched. In essence, the perturbation was a directional one and did not affect movement extent. There is strong evidence for independent planning and specification of movement direction and extent in the sensorimotor system (Favilla et al., 1989; Favilla et al., 1990; Fu et al. 1993; Gordon et al., 1994; Fu et al., 1995; Messier & Kalaska, 1997; Mutha and Sainburg 2007). This separation appears to extend right down to the neuronal level. For instance, Ebner and colleagues (Fu et al., 1993; Fu et al., 1995) showed that the while the same motor and premotor cortical neurons can encode direction, amplitude, and target position, these parameters are processed sequentially, giving rise to distinct neural representations. Correspondingly, more recent studies suggest that errors in direction and extent are also independently monitored and processed (Krakauer et al., 2000; Mutha et al., 2008; Shingane et al., 2025). Given these differences, the question of whether the findings of Kitchen et al. (2023), which employed directional perturbations, generalize to conditions where movement extent is perturbed, remains open.

Therefore, in the current study, we first asked whether perturbations that differentially alter the contribution of each arm to the extent of the shared cursor’s motion along the target direction axis, elicit flexible task-dependent responses as seen for directional perturbations. We predicted that varying the individual arm contributions to movement extent in a shared-cursor task would lead to differential arm responses, specifically resulting in asymmetric modulation of endpoint variability along movement extent. We found this to indeed be the case, while also replicating the results of Kitchen et al (2023) with directional perturbations applied in isolation. We next probed whether the sensorimotor system could resolve simultaneously imposed alterations along *both* movement extent and direction. If so, then altering the contribution of each arm simultaneously along these axes would lead to flexible task-dependent modulation of motor variability along both the extent and the direction axes. Alternatively, a unified response across both axes may emerge if the shared task goal constrains the independent resolution of errors along extent and direction when imposed simultaneously. We tested this idea in our next two experiments and again observed remarkable flexibility in the responses of the two limbs.

## MATERIALS AND METHODS

### Participants

Sixty right-handed subjects (44 male, aged between 18-29 years) participated in the study comprising of four experiments (n = 15 each). None of the participants reported any neurological or psychiatric disorders, had no joint or ligament disabilities and no other motor problems with the upper limbs. All participants gave written informed consent prior to participation. The study was approved by the Institute Ethics Committee of the Indian Institute of Technology Gandhinagar. Participants were monetarily compensated for their time.

### Setup

Participants sat in a height-adjustable chair facing a two-dimensional virtual reality set up, which consisted of a high-definition television display mounted horizontally over a tabletop (Figure 1A). A semi-silvered mirror, positioned between the table and the display, reflected visual feedback of hand position, along with start and target locations for the point-to-point reaching task that the participants would perform. The mirror configuration occluded direct arm vision, forcing participants to rely solely on the reflection displayed on the mirror. Participants performed bimanual arm movements (simultaneous movement of both arms) which were driven by the shoulder and the elbow; they wore splints that restricted wrist motion. Endpoint positions of both arms were recorded using 6 degree-of-freedom Flock of Birds electromagnetic sensors (Ascension trakSTAR) attached to the tip of the index fingers. Sensor position data was used to compute the velocity and acceleration of each arm.

**Figure 1:**
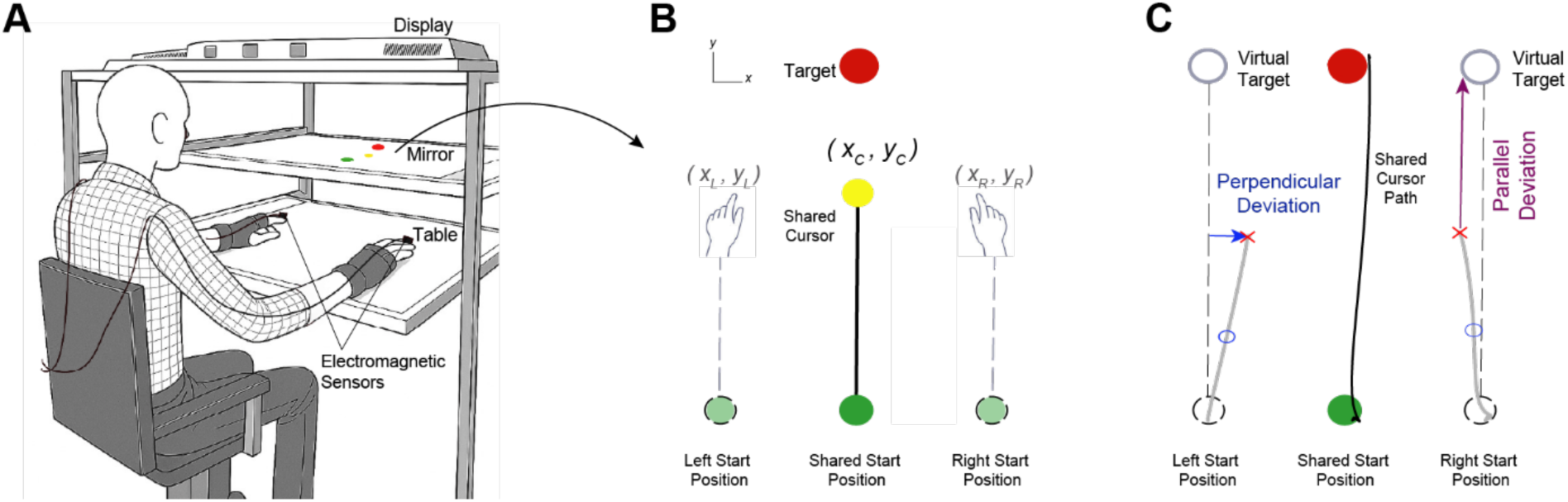
(A) Experimental setup with a horizontally mounted display and a mirror positioned above a tabletop. Participants performed bilateral reaching movements on the table while viewing arm position feedback on the mirror. Arm positions were tracked via electromagnetic sensors attached to the tip of the two index fingers. (B) Participants initiated each trial by aligning the left (*x_L_*, *y_L_*) and right (*x_R_*, *y_R_*) arm within separate start positions (dashed green circles), after which the display switched to a shared start position (solid green circle) and a single, shared feedback cursor ((*x_C_*, *y_C_*), yellow circle). Upon target (red circle) appearance, participants made rapid bimanual reaches to bring the shared cursor to the target. (C) Example arm and cursor trajectories from a single trial from the Equal_1 condition, where the gain ratios were changed to 2:2. The solid grey lines depict left and right arm trajectories; the black line represents the shared cursor path. Dashed grey lines indicate straight-line paths from each arm’s start position to its virtual target (unfilled grey circles), used to compute deviations. Virtual targets were not shown to the participants. Blue circles and red crosses denote arm positions at peak velocity and movement offset, respectively. The blue arrow indicates the perpendicular deviation at movement offset calculated as the lateral or medial deviation of each arm from the ideal trajectory. The violet arrow shows the parallel deviation at movement offset calculated as an undershoot or overshoot of each arm relative to the virtual target.

### Experimental Task

The main task involved making quick center-out bimanual reaching movements from a single, shared start position to a single target (red circle, 2.5 cm diameter). Each trial began with the presentation of two separate start positions (green circles, 1 cm diameter), one for each arm, positioned 10 cm to the left and right from the center of the workspace (Figure 1B). When participants brought the two arms within a 3 cm radius of each start circle, two feedback cursors (yellow circles, 0.8 cm diameter) were displayed on the screen to represent their hand position underneath, on the tabletop. To initiate a trial, participants had to align the two yellow cursors precisely within their respective start circles. Once they remained there for 500 ms, the individual starting positions and cursors were replaced by a shared starting position (green circle, 2 cm diameter) and a shared feedback cursor respectively. Subsequently, after a variable delay ranging from 450 to 1350 ms, a target appeared 16 cm ahead of the shared start circle with an audio beep signal, which served as the “go” cue. Participants were instructed to make a single, smooth straight movement to bring the shared cursor to the displayed target by moving both limbs together, as quickly as they could. At the end of each trial, they received a fixed reward of 10 points as feedback, regardless of their performance. The reward aimed to sustain motivation and minimise awareness of sudden change of visual feedback “gain” conditions across blocks (see “Visual feedback manipulation” below). Participants then moved their hand back to the start position, with the cursor hidden until they were within 3 cm of the start circle.

### Visual feedback manipulation

The relationship between the positions of the two arms on the tabletop and their feedback (shared yellow cursor) on the screen was manipulated in blocks across all four experiments. The position of the shared feedback cursor (*x_c_*, *y_c_*) was calculated in real-time based on the position of right arm (*x_R_*, *y_R_*), left arm (*x_L_*, *y_L_*) and gain parameters (GL_x_, GR_x_) and (GL_y_, GR_y_), as follows:

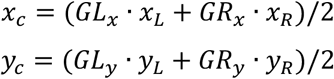

Here, GL_x_ and GR_x_ represented “perpendicular gain” parameters as they influenced the position of the cursor in a direction perpendicular to the direction of movement (i.e., the gains were applied to the x-coordinates of the left- and right-hand positions respectively). In contrast, GL_y_ and GR_y_ were gain parameters applied to the y-coordinates of the left- and right-hand positions respectively, and represented “parallel gain” since this manipulation occurred in (or in other words, parallel to) the direction of reach. By adjusting these parameters, we gave the two arms symmetric (equal contribution) or asymmetric (unequal contribution) control over the cursor’s perpendicular and parallel movements.

### Experimental Design

Each experiment comprised of two phases: familiarization and main task. During familiarization, participants completed 10 practice trials in which the perpendicular (GL_x_:GR_x_) and parallel (GL_y_:GR_y_) gain ratios of the left and right arm were fixed at 1:1. The main task involved 6 blocks of 50 trials each in the following order: No gain (1:1), Equal_1 (2:2), Asymmetric_1, Equal_2 (2:2), Asymmetric_2, Equal_3 (2:2). The manipulation employed in the two asymmetric blocks was counterbalanced across participants. All blocks, except for the two asymmetric blocks, were categorized as symmetric since the gain parameters applied during the hand-to-cursor transformation were identical for both arms. The No Gain block remained consistent across all four experiments, where visual feedback was identical to the familiarization phase. The values of gain parameters applied in each experiment are summarized in Table 1.

**Table 1:** Perpendicular and Parallel gain parameters applied to the left and right arm across 4 experiments.

| E<br>x | Equal blocks |  |  |  | LH High |  |  |  | RH High |  |  |  |
| --- | --- | --- | --- | --- | --- | --- | --- | --- | --- | --- | --- | --- |
|  | Perp Gain |  | Para Gain |  | Perp Gain |  | Para Gain |  | Perp Gain |  | Para Gain |  |
| P | GL<sub>x</sub> | GR<sub>x</sub> | GL<sub>y</sub> | GR<sub>y</sub> | GL<sub>x</sub> | GR<sub>x</sub> | GL<sub>y</sub> | GR<sub>y</sub> | GL<sub>x</sub> | GR<sub>x</sub> | GL<sub>y</sub> | GR<sub>y</sub> |
| 1 | 2 | 2 | 1 | 1 | 3.8 | 0.2 | 1 | 1 | 0.2 | 3.8 | 1 | 1 |
| 2 | 1 | 1 | 2 | 2 | 1 | 1 | 3.8 | 0.2 | 1 | 1 | 0.2 | 3.8 |
| 3 | 2 | 2 | 2 | 2 | 3.8 | 0.2 | 0.2 | 3.8* | 0.2 | 3.8 | 3.8 | 0.2** |
| 4 | 2 | 2 | 2 | 2 | 3.8 | 0.2 | 3.8 | 0.2<sup>#</sup> | 0.2 | 3.8 | 0.2 | 3.8<sup>##</sup> |

\* and \*\* are the gain parameters from Experiment 3 for the LH Perp–RH Para High and RH Perp–LH Para High conditions respectively. # and ## are the gain parameters for LH Perp–Para High and RH Perp–Para High from Experiment 4.

#### Experiment 1

The main task of the experiment 1 started with the No Gain block. In the subsequent blocks, we maintained the left and right parallel gain parameters at 1:1, while manipulating only the perpendicular gain (Kitchen et al., 2023). In the Equal_1 block, we increased the perpendicular gain ratio to 2:2 to smoothly enable the greater asymmetry that would appear in the ensuing asymmetric blocks. After the 2:2 block, participants completed the two asymmetric blocks (Asymmetric_1 and Asymmetric_2), labelled Left Hand Perpendicular High (LH Perp High) and Right Hand Perpendicular High (RH Perp High), interspersed with Equal_2 (2:2) and Equal_3 (2:2) blocks. In the “LH Perp High” block, the contribution of the left arm to the lateral (*x*-coordinate) motion of the shared cursor was increased (GL_x_ = 3.8), while the contribution of the right arm was decreased (GR_x_ = 0.2). Conversely, in the “RH Perp High” block, the perpendicular gain parameters were reversed (0.2:3.8), resulting in an increased contribution of the right arm and decreased the contribution of the left arm to lateral cursor motion.

#### Experiment 2

In experiment 2, the general structure of the blocks remained identical to experiment 1. However, to isolate the effects of gain manipulations along movement extent, we varied only the parallel gains while keeping the left and right perpendicular gains constant at 1:1 across all blocks. In the three Equal blocks, a uniform parallel gain of 2:2 was applied. The asymmetric blocks, labelled Left Hand Parallel High (LH Para High) and Right Hand Parallel High (RH Para High), had differential parallel gain parameters. In the “LH Para High” block, the contribution of the left arm to the parallel (y-coordinate) motion of the cursor was increased (GL_y_ = 3.8) while the contribution of the right arm was decreased (GR_y_ = 0.2). In the “RH Para High”, we altered the parallel gains to 0.2:3.8 to increase the contribution of the right arm and decrease the contribution of the left arm to the parallel motion of the cursor.

#### Experiment 3

The general procedure for our third experiment remained similar to the previous two experiments. However, the perpendicular and parallel gain parameters in the Equal gain blocks were maintained at 2:2 for the left and right arms. In the asymmetric gain blocks, we changed the contributions in the perpendicular and parallel directions simultaneously: one arm contributed more to perpendicular and less to parallel cursor motion, while the other had opposite contributions. The asymmetric blocks were labelled LH Perp–RH Para High and RH Perp–LH Para High. In the “LH Perp–RH Para High” block, the perpendicular gains were set at 3.8:0.2 while the parallel gains were 0.2:3.8. These gains allowed us to introduce a higher contribution of left arm to the lateral motion and lower contribution to the parallel motion of the shared cursor. On the contrary, the right arm contributed less to the perpendicular motion and more to the parallel motion of the cursor. This was reversed in the “RH Perp–LH Para High” block, with perpendicular gain parameters set at 0.2:3.8 and parallel gain parameters set at 3.8:0.2. Thus, in this condition, the right arm had increased lateral but decreased parallel contribution, while the left arm had increased parallel but decreased lateral contribution to the motion of the cursor.

#### Experiment 4

In our fourth experiment, the same arm simultaneously contributed more to both perpendicular and parallel cursor motion, while the other arm contributed less in both dimensions. Again, the perpendicular and parallel gain parameters of left and right hand in the Equal blocks were 2:2. Two asymmetric blocks were introduced: LH Perp–Para High and RH Perp–Para High. In the “LH Perp–Para High” block, the perpendicular and parallel gain ratios were both set at 3.8:0.2, which increased the contribution of the left arm and decreased the contribution of the right arm to both perpendicular and parallel motion of the cursor. Conversely, in the “RH Perp–Para High” block, the gain ratios were reversed to 0.2:3.8, resulting in increased contribution of the right arm and decreased contribution of the left arm to perpendicular as well as parallel motion of the cursor.

### Data Analysis

The hand position data obtained from the trakStar sensors were low-pass Butterworth filtered with a cutoff frequency of 8 Hz. Data from the familiarization phase were not analyzed. Rather, we focused on the *x* and *y* positions of the two arms in the main task (movements were constrained largely to the horizontal plane). Tangential velocity profiles for each arm were derived by differentiating their positions over time. Movement onset for each arm was defined as the first timepoint where movement velocity fell below 8% of the peak velocity, identified by tracing backwards from the point of peak velocity. Movement offset was similarly defined at the first point after peak velocity where the velocity fell below the same 8% threshold. To time-lock the movement offset and peak velocity of the left and right arms for our covariation analysis, we averaged the respective time point of each measure across the two arms and defined a “new” movement offset and peak velocity based on this averaged time point (Kitchen et al., 2023). Trials with no or incomplete movement, identified using movement onset and offset, were excluded from the analysis (0.38% of all trials). To assess performance in the task, we computed endpoint shared cursor error along each axis (perpendicular and parallel) as the difference between the final position of the cursor at movement offset and the corresponding target coordinate (*x* or *y*). To assess differences across gain conditions in mean cursor errors, we used a Kruskal–Wallis test within each cycle (10 trial epochs), followed by Dunn’s post hoc test for pairwise comparisons when significant effects were found.

Additionally, two measures of deviations in hand trajectory, perpendicular deviation and parallel deviation, were computed for each arm. Furthermore, this computation was performed at two timepoints: the point of peak velocity and movement offset. Perpendicular deviation was defined as the displacement of *x*-coordinate of the arm (at peak velocity or at movement offset) from the straight-line joining its start position to its corresponding “virtual” target (Figure 1C). This captured lateral and medial deviations of the arm from the “ideal” straight trajectory. Parallel deviation was defined as displacement of the *y*-coordinate of the arm (at peak velocity or at movement offset) from the *y*-coordinate of the virtual target; this represented movement undershoot or overshoot. Note that the virtual targets were not displayed to the participants, but were defined only for analysis purposes as essentially the locations that the two arms would have to reach had the perpendicular and parallel gains been maintained at 1:1.

#### Covariance analysis

To assess changes in the coordination of movement direction, we computed the covariance between the left and right arm perpendicular deviations across each block. Similarly, to quantify changes along movement extent, we calculated the covariance between the parallel deviations of the left and right arms. This was done by fitting Deming orthogonal regression models with equal variance to individual participants and to group-level deviations of the two arms (data from all subjects combined). We used orthogonal regression to account for measurement errors in both left and right arm deviations. The raw slopes obtained from the regression analysis were converted to slopes angles in degrees. Notably, for perpendicular deviations, we subtracted 180° from the slope angles ranging between +45° to +90° to preserve the continuous decreasing nature of the slope while crossing the *y*-axis. No such transformations were required for the slopes of the regression of parallel deviations because the slopes decreased to zero at the *x*-axis and continued to increase beyond it.

Fitting orthogonal regression to individual participants data resulted in highly variable slopes, likely due to the limited number of trials per block (50 trials), and therefore, fitting orthogonal regression to the group data was considered more reliable (Kitchen et al., 2023). The slope of the orthogonal fit was compared between Equal_1, Asymmetric_1 and Asymmetric_2 blocks to reveal changes in coordination patterns. A slope of 45° or –45° indicated strong positive or negative covariation respectively, implying an equal change in the deviation of the two arms. In contrast, slopes deviating from these values reflected asymmetry in the response of the two arms. Further, positive covariation indicated that the two arms deviated in the same direction (anti-symmetric responses in the two limbs for perpendicular deviations, and both arms either undershooting or overshooting for parallel deviations), whereas negative covariation suggested that the arms deviated in opposite directions (symmetric responses in the two limbs for perpendicular deviations, and one arm overshooting and the other undershooting in case of parallel deviations). Thereafter, 95% confidence ellipses were computed for individual and group slopes. In addition to these analyses, we used linear mixed models to analyse the effect of asymmetric gain change on individual slopes. The Equal_1 condition and the two asymmetric conditions were modelled as a continuous fixed-effect predictor calculated as the ratio of right-hand gain parameter to the total gain. For instance, in the RH Para High with a gain ratio of 0.2:3.8 (GL_y_: GR_y_), the fixed-effect value was 0.95 reflecting the relative contribution of the right hand to the shared cursor movement in that direction. The continuous fixed factor allowed us to assess how increasing right-hand contribution to the shared cursor influenced orthogonal slope estimates. The model included random intercepts and slopes for each participant to account for individual variability.

#### Bootstrapping

To statistically confirm the adaptive changes in coordination observed at the group-level in the two asymmetric blocks, we performed a bootstrapping analysis. We took the original 15 participants and created a new group of 15 randomly selected participants while allowing the same participant to appear multiple times (that is, with replacement). This resampling was repeated 10000 times. Each resampled dataset represented a new “group”, on which we applied orthogonal regression models to the group-level perpendicular and parallel deviations of the two arms, generating a distribution of 10000 regression slopes. This was done for the two asymmetric blocks. We used this method to address the issue of variability in slopes across individual participants, and account for potentially extreme slopes (while being non-outliers) relative to the group (Hardwick et al., 2017). By allowing participants to be selected multiple times or not at all in each resample, we accounted for the inherent heterogeneity within the participant population. The 95% confidence intervals for the bootstrapped distribution were obtained by using bias-corrected and accelerated bootstrap (BCa) method (D’Cruz et al., 2024; Puth et al., 2015) to account for the skewness and bias in the distribution. These confidence intervals were then compared across the asymmetric gain conditions.

## RESULTS

### Experiment 1

In our first experiment, we attempted to replicate the findings of Kitchen et al. (2023); we increased the contribution of one arm to the perpendicular motion of the cursor while reducing the contribution of the other arm, in a blocked manner. Each block represented one of three conditions for the contribution of left and right arm to the *x*-coordinate of the cursor: 1) RH Perp High, where the right arm contributed more while the left arm contributed less, 2) Equal, where both arms contributed equally, and 3) LH Perp High, in which the left arm contributed more while the right arm contributed less. In line with Kitchen et al. (2023), we predicted that the sensorimotor system would reduce the medial-lateral variability of the arm with higher contribution to the cursor motion while allowing the other arm to vary more freely.

We first analyzed the mean endpoint cursor errors along the *x*-axis (Figure 2A) across cycles (10 trials epochs) for the Equal_1, LH Perp High and RH Perp High conditions. Mean errors were negative for LH Perp High and positive for RH Perp High, reflecting direction-specific effects of the gain manipulation. A Kruskal-Wallis test revealed a significant effect of gain condition in all cycles (all p < 0.002, Table 2) except for cycles 3 (p = 0.2235) and 5 (p = 0.0595). Post-hoc Dunn’s test showed significant differences between LH Perp High and RH Perp High conditions in cycle 1 (p = 0.036), cycle 2 (p = 0.0013), and cycle 4 (p = 0.0080), but other comparisons did not reach statistical significance.

**Figure 2:**
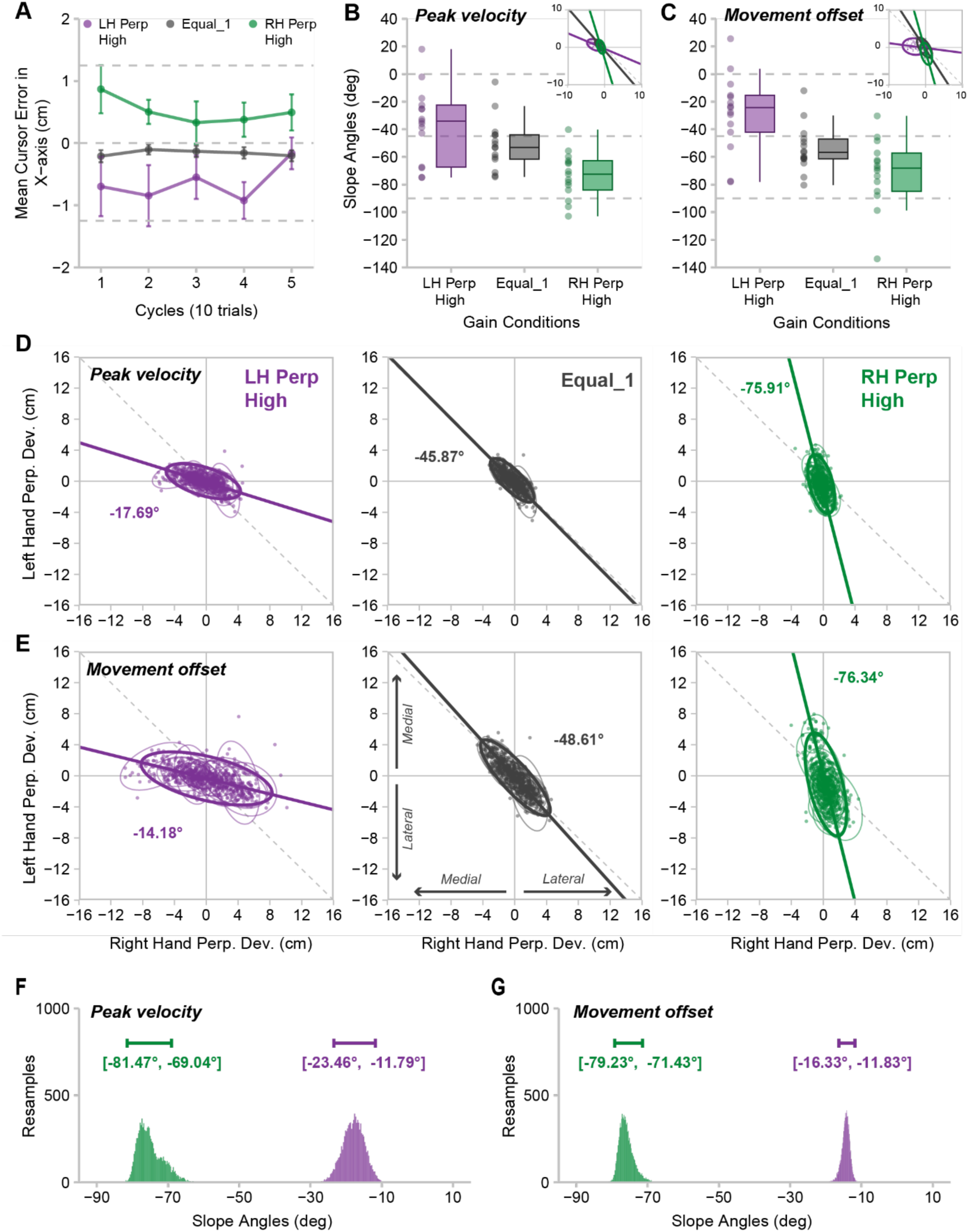
(A) Mean±SEM shared cursor error along the x-axis at movement offset across five cycles plotted for the three gain conditions: LH Perp High (purple), Equal_1 (grey), and RH Perp High (green). Positive values indicate rightward errors with respect to the center of the target; negative values indicate leftward errors. Dashed lines at ±1.25 cm mark the lateral boundaries of the target. **(B, C)** Boxplots of regression slopes computed for the perpendicular deviations of the right and left arms at **(B)** peak velocity and **(C)** movement offset for the three gain conditions. Lines at 0°, –45°, and –90° denote theoretically predicted slope angles for the perpendicular deviations of LH Perp High, Equal_1 and RH Perp High conditions respectively. Dots represent individual participants. **(INSET)** Covariance plots of perpendicular deviations for a representative participant, overlaid across conditions at **(B)** peak velocity and **(C)** movement offset. **(D, E)** Covariance between perpendicular deviations of the right and left arms at **(D)** peak velocity and **(E)** movement offset for the three gain conditions. Each panel shows data from all trials across all participants. Darker ellipses denote 95% confidence ellipses of group-level data, and the dark lines indicate group-level orthogonal regression fits (annotated values). Lighter ellipses reflect confidence ellipses for individual participants. Dashed line (slope = –45°) denotes the line of equivalence. Positive deviations in the left arm and negative deviations in the right arm represent medial perpendicular deviations. **(F, G)** Bootstrapped distributions (10,000 resamples) of the overall, group-level orthogonal regression slope for the two asymmetric gain conditions (LH Perp High: purple; RH Perp High: green), at **(F)** peak velocity and **(G)** movement offset. Bias-corrected and accelerated (BCa) 95% confidence intervals are presented and annotated above each distribution. Non-overlapping intervals indicate statistically distinct coordination strategies between the two conditions.

**Table 2:**
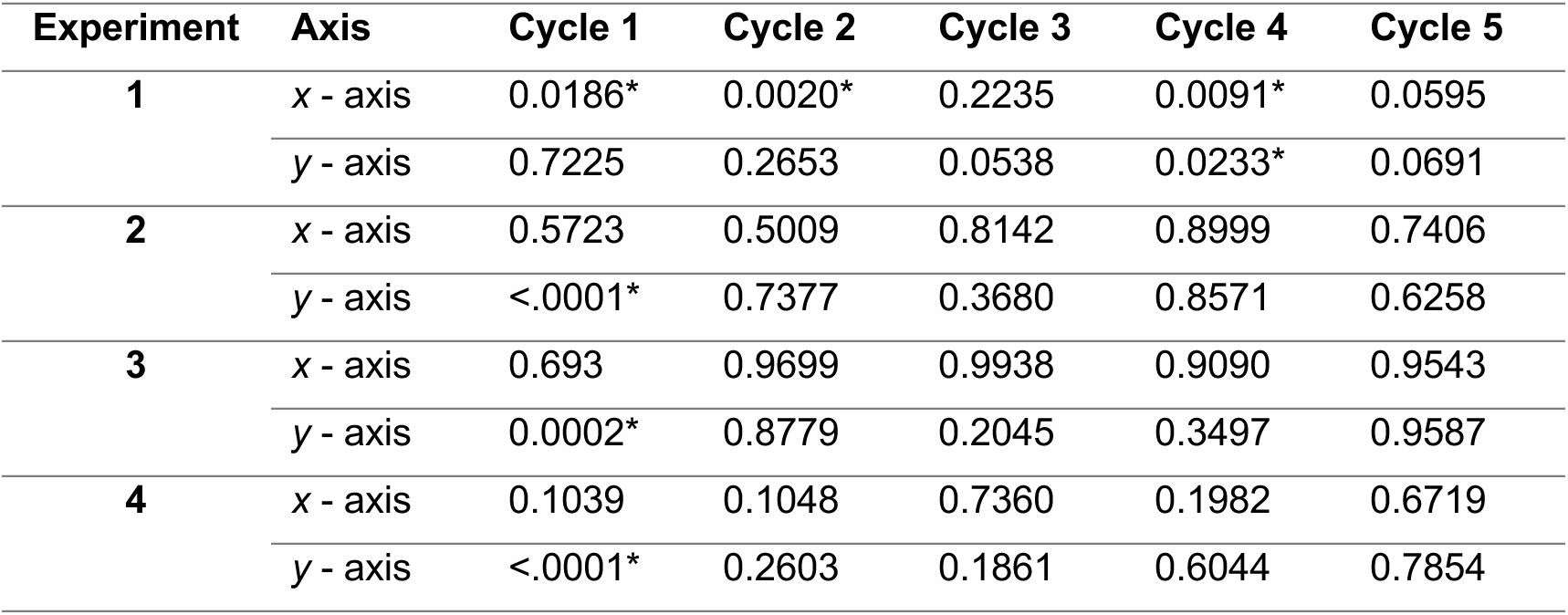
P-values from Kruskal-Wallis tests comparing cursor errors across gain conditions (Equal_1, Asymmetric_1, Asymmetric_2) for each cycle (10-trial epochs) in both the x- and y-axis directions, reported separately for all four experiments. Statistically significant differences are indicated by an *.

To investigate changes in coordination between the two arms following the introduction of the gains, we analyzed the covariance between the perpendicular deviations of the right and left arms at peak velocity and movement offset across the Equal_1, LH Perp High, and RH Perp High conditions. The degree of covariance was quantified using Deming regression slopes, obtained by fitting an orthogonal linear regression to the perpendicular deviations of the left arm as a function of the right arm. Figures 2B and 2C show the regression slopes for individual participants at peak velocity and movement offset, respectively. Insets in each panel illustrate how these slopes changed for a single participant across the gain conditions. In the Equal_1 condition, where both arms contributed equally to the motion of the shared cursor, the slope was close to the line of equivalence (slope = −45°) indicating a strong negative covariation between the perpendicular deviations of the two arms. This suggested that the medially-directed perpendicular response of one arm was compensated by an almost equal medially-directed response of the other arm, and vice versa, in order to hit the target accurately.

A change in slope from that in the Equal_1 condition in the asymmetric LH Perp High and RH Perp High conditions would provide evidence favouring dynamic, task-dependent modulation of this coordination pattern. Consistent with Kitchen et al. (2023), we found that for the LH Perp High condition, where the left arm contributed more to the perpendicular motion of the cursor and the right arm contributed less, the slope shifted away from the line of equivalence, indicating weakened covariance between the arms. Likewise, in the RH Perp High condition, where the right arm contributed more to the perpendicular motion of the cursor, the regression slope shifted in the opposite direction. This trend was confirmed at the group level using a linear mixed model fit on the regression slopes obtained from individual participants for the three gain conditions. We found a significant effect of gain conditions on the regression slopes, both at peak velocity (Fixed effect slope = −41.22°, 95% CI = [−60.27°, −22.17°], p < 0.001, Figure 2B) and at movement offset (Fixed effect slope = −47.90°, 95% CI = [−67.88°, −27.92°], p < 0.001, Figure 2C).

Figure 2D shows the covariance relationships between right- and left-hand perpendicular deviations at peak velocity, with all trials from all participants plotted on the same axes and with 95% confidence ellipses computed for individual (lighter ellipses) and group data (darker ellipses). Task-dependent modulations of the covariance between the perpendicular deviations of the two arms were already evident at this time. The group regression slope was −45.87° in the Equal_1 condition, again pointing to a strong negative covariation between the arms. In the LH Perp High condition, the slope changed to −17.69°, suggesting that this strong covariation had been weakened due to the gain change. The confidence ellipse (Figure 2D, dark purple ellipse) revealed restricted spatial variance for the left arm compared to the right. In contrast, in the RH Perp High condition, the slope changed to −75.91°, with the confidence ellipse (Figure 2D, dark green ellipse) now pointing to restricted variance for the right arm with the left arm being allowed to vary more freely. The same pattern was evident at movement offset (Figure 2E). In the Equal_1 gain condition, the group regression slope was −48.61°. In the LH Perp High condition, the slope changed to −14.18°, while in the RH Perp High condition, it was modulated to −76.34°. Notably, the changes in covariance in the two asymmetric conditions emerged rapidly: group-level slopes showed clear divergence relative to the Equal_1 condition within the first 10 trials and continued to evolve throughout the block (See exp. 1 Supplementary Results, Figure S1). The covariance of perpendicular deviations also adapted dynamically, responding not only to the main experimental blocks but also to the interleaved Equal blocks. Specifically, while covariance weakened in the asymmetric conditions, the regression slopes (at movement offset) realigned with the line of equivalence, during the interleaved Equal_2 (Slope = −48.12°) and Equal_3 (Slope = −43.25°) conditions. This highlighted the ability of the sensorimotor system to adopt a flexible coordination strategy. To compare the coordination patterns observed at the group-level, we used a bootstrapping analysis. We resampled with replacement, fit an orthogonal regression model for each resample and generated a distribution of 10000 slopes for each condition at peak velocity (Figure 2F) and movement offset (Figure 2G). At peak velocity, we determined the 95% confidence intervals from these distributions as [−23.46°, −11.79°] for the LH Perp High and [−81.47°, −69.04°] for RH Perp High conditions. At movement offset, the corresponding intervals were [−16.33°, −11.83°] for LH Perp High and [−79.23°, −71.43°] for RH Perp High conditions. The non-overlapping confidence intervals between the two asymmetric conditions at both these time points indicated statistically significant differences, suggesting clearly distinct coordination strategies between these conditions.

Next, we examined whether the flexibility in bimanual coordination observed in the perpendicular deviations also occurred in the forward reaching (parallel) direction in this experiment where only perpendicular gains were manipulated. We first analyzed cursor errors along the *y*-axis across the three gain conditions. Mean errors were consistently positive for all the three gain conditions (Figure 3A). A Kruskal-Wallis test revealed no significant effects of the perpendicular gain on the mean cursor errors in all cycles (all p > 0.054, Table 2), except for cycle 4 (p = 0.023); post-hoc comparisons showed a significant difference only between the LH Perp High and Equal_1 conditions (p = 0.024). Next, we used a linear mixed model to assess whether regression slopes differed across the three gain conditions. We found no significant effect of gain condition either at peak velocity (Fixed effect slope = 0.13°, CI = [−3.02°, 3.28°], p = 0.931, Figure 3B) or at movement offset (Fixed effect slope = 1.32°, 95% CI = [−4.85°, 7.48°], p = 0.65, Figure 3C).

**Figure 3:**
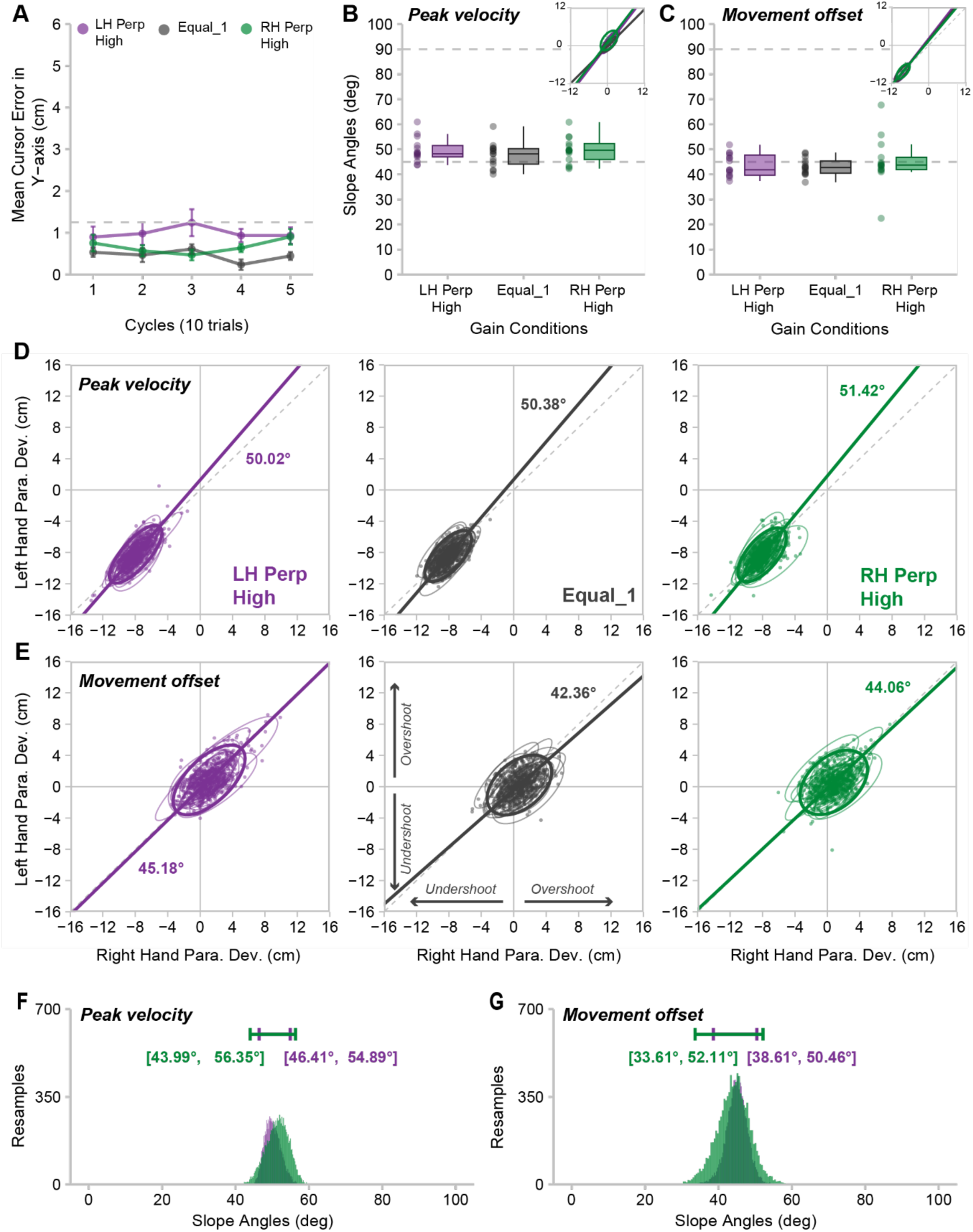
(A) Mean±SEM shared cursor error along the y-axis at movement offset, across five cycles, for the three gain conditions: LH Perp High (purple), Equal_1 (grey), and RH Perp High (green). Positive values indicate overshoot while negative values indicate undershoot of the shared cursor with respect to the target center. Dashed line at 1.25 cm indicates the upper vertical boundary of the target. **(B, C)** Boxplots of regression slopes computed for the parallel deviations of the right and left arms at **(B)** peak velocity and **(C)** movement offset for the three gain conditions. Dots represent individual participants. **(INSET)** Covariance plots of parallel deviations for a representative participant, overlaid across conditions at **(B)** peak velocity and **(C)** movement offset. **(D, E)** Covariance between parallel deviations of the right and left arms at **(D)** peak velocity and **(E)** movement offset for the three gain conditions. Each panel shows data from all trials across all participants. Darker ellipses denote 95% confidence ellipses of group-level data, and the dark lines indicate group-level orthogonal regression fits (annotated values). Lighter ellipses reflect confidence ellipses for individual participants. Dashed line (slope = 45°) denotes the line of equivalence. Positive deviations indicate overshoot, and negative deviations indicate undershoot relative to the virtual target. **(F, G)** Bootstrapped distributions (10,000 resamples) of the overall, group-level orthogonal regression slope for the two asymmetric gain conditions (LH Perp High: purple; RH Perp High: green), at **(F)** peak velocity and **(G)** movement offset. Bias-corrected and accelerated (BCa) 95% confidence intervals are presented and annotated above each distribution. Non-overlapping intervals indicate statistically distinct coordination strategies between the two conditions.

The covariance relationships between the parallel deviations of the two arms, at peak velocity and movement offset, revealed similar findings. At peak velocity (Figure 3D), the Equal_1 condition (Slope = 50.38°) revealed a positive covariation, indicating that when one arm overshot or undershot its virtual target, the other arm tended to do the same. Notably, there was not much of a change in the regression slopes for parallel deviations in the LH Perp High (Slope = 50.02°) and RH Perp High (Slope = 51.42°) conditions. A similar pattern was observed at movement offset (Figure 3E), where the regression slopes across all three conditions remained comparable. Further, bootstrap analyses confirmed these results with overlapping confidence intervals at peak velocity (Figure 3F) for the LH Perp High (CI = [46.41°, 54.89°]) and RH Perp High (CI = [43.99°, 56.35°]) conditions, as well as at movement offset (LH Perp High: CI = [38.61°, 50.46°]; RH Perp High: CI = [33.61°, 52.11°], Figure 3G). These analyses reinforced the findings that perpendicular gain manipulations did not modulate coordination along the movement extent axis.

### Experiment 2

The findings from experiment 1 demonstrated that bimanual coordination is sensitive to contributions of each arm to the perpendicular motion of a common, shared cursor. In experiment 2, we investigated whether a similar adaptive and task-dependent mechanism is employed when the arm contributions to the shared cursor motion are asymmetrically manipulated along the target direction axis. In the LH Para High and RH Para High conditions, the left and right arms respectively, had increased contributions to the cursor’s parallel motion. To isolate the impact of these parallel gain variations, perpendicular gains were held constant at 1:1 across all blocks. We expected that the regression slopes in the LH Para High and RH Para High blocks would diverge from those in the Equal_1 block; specifically, the arm with higher parallel contribution to the cursor would demonstrate reduced variability in extent.

In the No Gain (1:1) condition, participants made bimanual reaching movements of 15.44 cm and 15.05 cm magnitude with their right and left arms respectively, with a corresponding mean endpoint shared cursor error of 0.75 cm in the *y*-direction. In the following Equal_1 block, where the parallel gain was increased to 2:2, the mean shared cursor error along y-axis increased to 5.46 cm in the first 10 trials (Figure 4A). Analysis of arm trajectories from a representative participant during the Equal_1 block (inset Figure 4A) revealed a decrease in the mean reaching amplitude of the right and left arms to 8.44 cm and 8.71 cm respectively. The corresponding group-averaged movement magnitude was 9.29 cm for the right arm and 9.26 cm for the left arm, which indicated a uniform compensatory reduction in movement extent to minimize the shared cursor error. Cursor errors along the *y*-axis in the Equal_1 block reduced over time, with the mean error decreasing to 1.35 cm in the last 10 trials (Figure 4A). We conducted a Kruskal-Wallis test on cursor errors comparing across gain conditions in each 10-trial cycle (Table 2). Only cycle 1 showed a significant effect (p < 0.0001); post-hoc Dunn’s tests revealed that both LH Para High (p < 0.0003) and RH Para High (p < 0.0012) differed significantly from Equal_1, but not from each other (p = 1.00).

**Figure 4:**
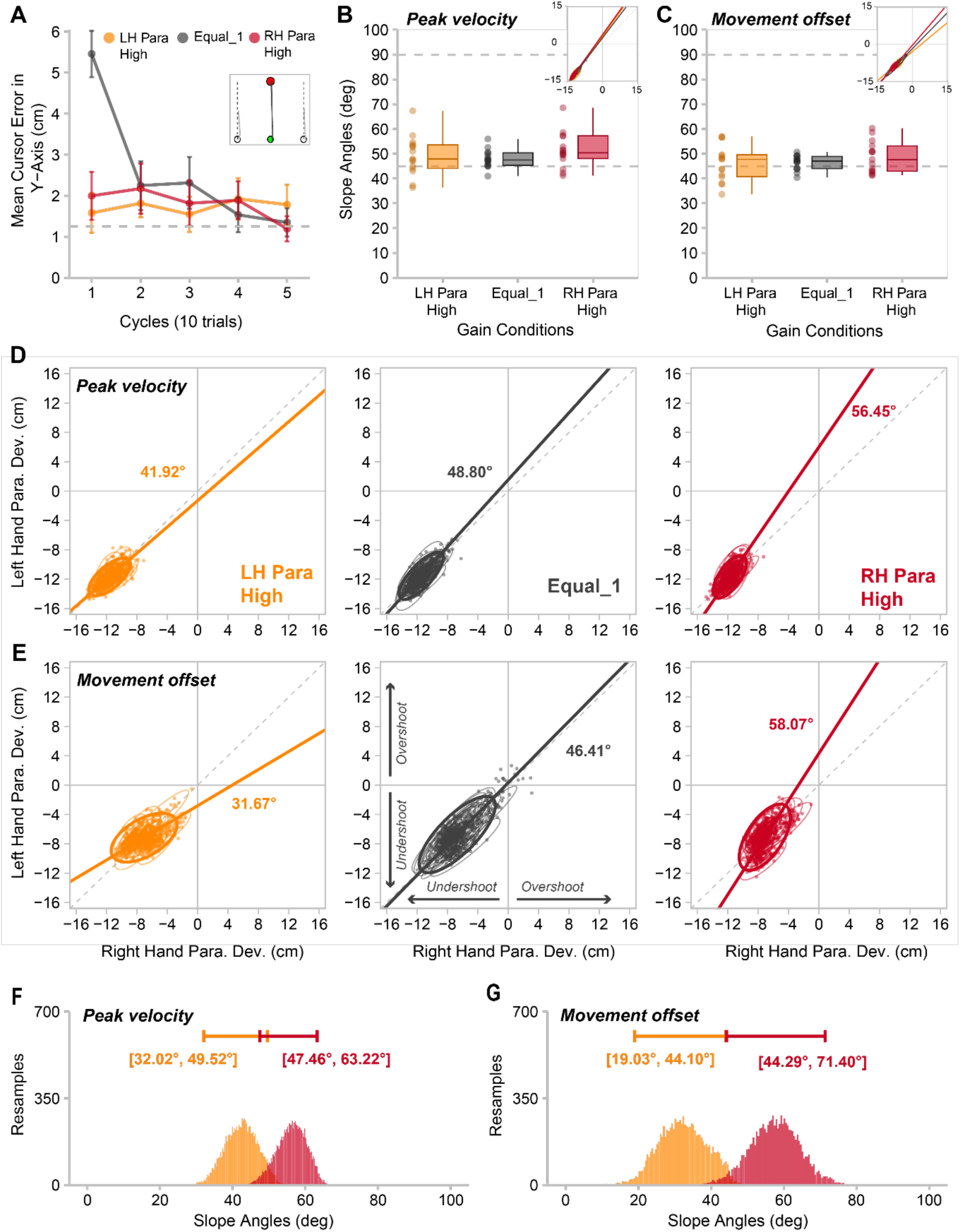
(A) Mean±SEM shared cursor error along the y-axis at movement offset, across five cycles, for the three gain conditions: LH Para High (orange), Equal_1 (grey), and RH Para High (red). Positive values indicate overshoot while negative values indicate undershoot of the shared cursor with respect to the center of the target. Dashed line at 1.25 cm indicates upper vertical boundary of the target. **(INSET)** Representative arm and cursor trajectories from a single trial from the Equal_1 condition where the gain ratios were changed to 2:2. Solid grey lines depict left and right arm trajectories; the black line represents the shared cursor path. **(B, C)** Boxplots of regression slopes computed for the parallel deviations of the right and left arms at **(B)** peak velocity and **(C)** movement offset for the three conditions. Lines at 0°, 45°, and 90° denote hypothesized slope angles for LH Para High, Equal_1 and RH Para High conditions respectively. Dots represent individual participants. **INSET** Covariance plots of parallel deviations for a representative participant, overlaid for across conditions at **(B)** peak velocity and **(C)** movement offset. **(D, E)** Covariance between parallel deviations of the right and left arms at **(D)** peak velocity and **(E)** movement offset for the three gain conditions. Each panel shows data from all trials across all participants. Darker ellipses denote 95% confidence ellipses of group-level data, and the dark lines indicate group-level orthogonal regression fits (annotated values). Lighter ellipses reflect confidence ellipses for individual participants. Dashed line (slope = 45°) denotes the line of equivalence. Positive deviations indicate overshoot, and negative deviations indicate undershoot relative to the virtual target. **(F, G)** Bootstrapped distributions (10,000 resamples) of the overall, group-level orthogonal regression slope for the two asymmetric gain conditions (LH Para High: orange; RH Para High: red), estimated at **(F)** peak velocity and **(G)** movement offset. Bias-corrected and accelerated (BCa) 95% confidence intervals are presented and annotated above each distribution. Non-overlapping intervals indicate statistically distinct coordination strategies between the two conditions.

To characterize the coordination between the two arms, we examined the covariance relationships between the parallel deviations of the right and left hands across the Equal_1, LH Para High, and RH Para High blocks. The insets of Figure 4B and 4C show the changes in regression slope across conditions for a single participant at peak velocity and movement offset. At peak velocity, the regression slopes for Equal_1, LH Para High and RH Para High were generally aligned, indicating similar covariation across conditions, but their deviation from the line of equivalence reflected a generally weakened covariance. At movement offset, both the conditions diverged away from Equal_1 in opposite directions, revealing task-dependent modulation. However, an analysis of the regression-slopes obtained from individual participants using a linear mixed-effect model revealed no significant effect of gain conditions at either peak velocity (Fixed effect slope = 2.52°, 95% CI = [−0.76°, 5.79°], p = 0.125, Figure 4B) or movement offset (Fixed effect slope = 2.69°, 95% CI = [−1.39°, 6.77°], p = 0.179, Figure 4C).

Figure 4D shows the group-level covariance relationships between the parallel deviations of right and left arms at peak velocity. The slope in the Equal_1 condition, where both arms contributed equally to the parallel motion of the cursor, was 48.80°, indicating strong positive covariation between the parallel deviations of the two arms. Task-dependent modulation at this stage of movement was evident only for the RH Para High condition (56.45°), but not in the LH Para High condition (41.92°), suggesting that changes in coordination occurred early when the right arm had higher contribution to the parallel motion of the shared cursor.

However, a more robust change in coordination was observed for both arms at movement offset (Figure 4E). In the Equal_1 block the regression slope was 46.41°, with the group confidence ellipse located entirely within the third quadrant. This indicated that both arms consistently moved less than before, which was also captured by the shift in the origin of the ellipse from (0.56, 0.95) in the No Gain block to (−6.71, −6.74) in the Equal_1 block. In the subsequent asymmetric conditions, where the contribution of one arm to the parallel motion of the cursor was increased while that of the other arm was reduced, the regression slopes shifted markedly from the Equal_1 condition. In the LH Para High condition, where the contribution of the left arm to the parallel motion of the cursor was increased and the right arm’s contribution was decreased, the regression slope changed to 31.67°, indicating a weakening of the covariation between the two arms in the parallel direction (Figure 4E). In contrast, in the RH Para High condition, where the right arm had higher contributions to the parallel motion of the cursor and the left arm had low contributions, the slope increased to 58.07°. These changes in covariance of parallel deviations in response to the asymmetric parallel gains emerged rapidly, within the first 10 – 20 trials of the block for both asymmetric conditions (See exp. 2 Supplementary Results, Figure S2). The regression slopes at movement offset gradually returned to values comparable to the Equal_1 block; we observed slopes of 53.17° in the Equal_2 and 46.22° in the Equal_3 conditions.

To account for the high variability in individual slopes, we bootstrapped the group-level slopes at peak velocity and movement offset, and compared them across the asymmetric conditions. At peak velocity (Figure 4F), the 95% confidence intervals partially overlapped between the LH Para High (CI = [32.02°, 49.52°]) and RH Para High (CI = [47.46°, 63.22°]) conditions, which suggested that distinct coordination strategies had not yet fully emerged at this stage of the movement. In contrast, at movement offset (Figure 4G), we found no overlap in the confidence intervals of the LH Para High (CI = [19.03°, 44.10°]) and RH Para High (CI = [44.29°, 71.40°]) conditions, pointing to statistically significant differences between the two. This indicated that participants employed distinct coordination strategies to account for the differential gains in these two conditions.

We then examined whether manipulating arm contributions to parallel motion of the cursor affected the flexibility in coordination in the perpendicular direction. A Kruskal-Wallis test revealed no significant differences in cursor errors along the *x*-axis (Figure 5A) across conditions in any cycle (all p > 0.50, Table 2). Further, a linear mixed-effects analysis on the regression slopes obtained from individual participants to probe for changes in covariance revealed no significant effect of gain conditions (at peak velocity: Fixed effect slope = 2.28°, CI = [−12.26°, 16.82°], p = 0.742, Figure 5B; at movement offset: Fixed effect slope = 4.41°, 95% CI = [−9.27°, 18.10°], p = 0.501, Figure 5C). We then examined group-level covariance relationships, shown in Figures 5D (at peak velocity) and 5E (at movement offset). At peak velocity (Figure 5D), the Equal_1 condition (−45.99°) revealed a strong negative covariation. Notably, the regression slopes for perpendicular deviations in the LH Para High (−40.68°) and RH Para High (−50.03°) conditions were not very different relative to the Equal_1 condition. Similarly, at movement offset (Figure 5E), no notable changes in group-level regression slopes across the three gain conditions were observed. This was also confirmed via our bootstrapping analysis, which revealed overlapping confidence intervals for the two asymmetric conditions both at peak velocity (LH Para High: [−56.99°, −29.67°]; RH Para High: [−64.83°, −33.38°], Figure 5F) and at movement offset (LH Para High: [−49.61°, −35.71°]; RH Para High: [−56.74°, −41.29°], Figure 5G), suggesting no task-dependent differences in perpendicular deviations across conditions.

**Figure 5:**
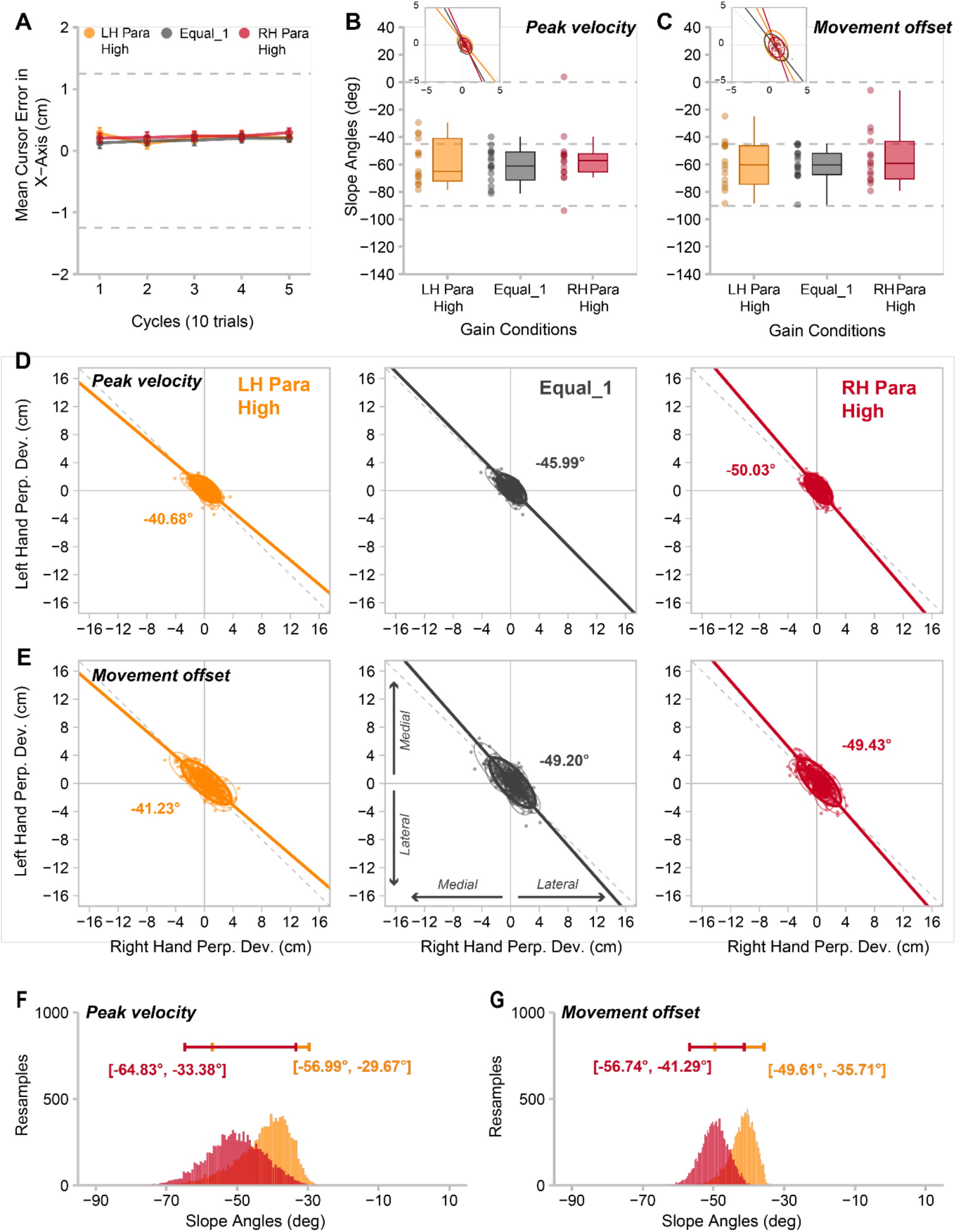
(A) Mean±SEM shared cursor error along the x-axis at movement offset across five cycles plotted for the three gain conditions: LH Para High (orange), Equal_1 (grey), and RH Para High (red). Positive values indicate rightward errors with respect to the center of the target; negative values indicate leftward errors. Dashed lines at ±1.25cm mark the lateral boundaries of the target. (B, C) Boxplots of regression slopes computed for the perpendicular deviations of the right and left arms at (B) peak velocity and (C) movement offset for the three gain conditions. Dots represent individual participants. (INSET) Covariance plots of perpendicular deviations for a representative participant, overlaid across conditions at (B) peak velocity and (C) movement offset. (D, E) Covariance between perpendicular deviations of the right and left arms at (D) peak velocity and (E) movement offset for the three gain conditions. Each panel shows data from all trials across all participants. Darker ellipses denote 95% confidence ellipses of group-level data, and dark lines indicate group-level orthogonal regression fits (annotated values). Lighter ellipses reflect confidence ellipses for individual participants. Dashed line (slope = –45°) denotes the line of equivalence. Positive deviations in the left arm and negative deviations in the right arm represent medial perpendicular deviations. (F, G) Bootstrapped distributions (10,000 resamples) of the group-level orthogonal regression slope for the two asymmetric gain conditions (LH Para High: orange; RH Para High: red), estimated at (F) peak velocity and (G) movement offset. Bias-corrected and accelerated (BCa) 95% confidence intervals are presented and annotated above each distribution. Non-overlapping intervals indicate statistically distinct coordination strategies between the two conditions.

### Experiment 3

Our first two experiments demonstrated decoupling of coordination between the arms that primarily emerged along the axis of gain manipulation. Notably, this modulation occurred irrespective of whether gain was applied to movement direction or extent. Building on these findings, we probed, in experiment 3, whether task-dependent coordination strategies persist when asymmetric perturbations are simultaneously imposed along both movement extent and direction. In one condition (LH Perp–RH Para High), the left arm contributed more to the perpendicular motion and less to the parallel motion of the cursor, whereas the right arm contributed more to the parallel motion while contributing less to its perpendicular motion. Conversely, in the RH Perp–LH Para High condition, the right arm had a higher contribution to perpendicular motion of the cursor while contributing less to its parallel motion, whereas the left arm had a higher contribution to its parallel motion while contributing less to its perpendicular motion. We expected that regression slopes would be modulated for both perpendicular and parallel deviations for the asymmetric conditions in correspondence with the specific nature of the perturbation.

Figure 6A shows the cursor errors along the *x*-axis across 10-trial cycles. In the first cycle, the LH Perp–RH Para High condition showed positive errors, while the RH Perp–LH Perp High condition had negative errors, reversing the pattern observed in experiment 1. A Kruskal-Wallis test showed no significant differences in cursor errors along the *x*-axis across gain conditions in any cycle (all p > 0.69, Table 2).

**Figure 6:**
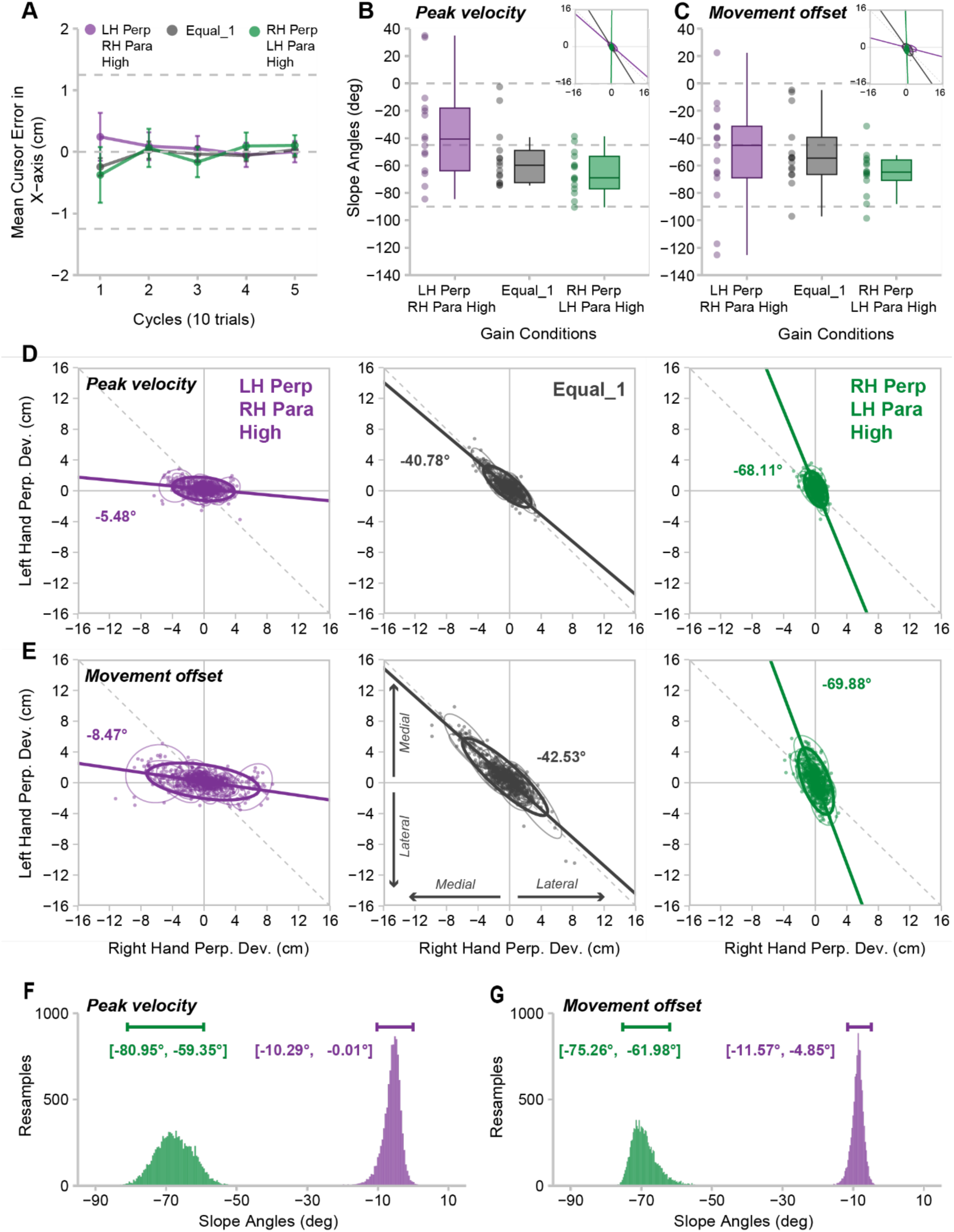
(A) Mean±SEM shared cursor error along the x-axis at movement offset across five cycles plotted for the three gain conditions: LH Perp–RH Para High (purple), Equal_1 (grey), and RH Perp–LH Para High (green). Positive values indicate rightward errors with respect to the center of the target; negative values indicate leftward errors. Dashed lines at ±1.25 cm mark the lateral boundaries of the target. **(B, C)** Boxplots of regression slopes computed for the perpendicular deviations of the right and left arms at **(B)** peak velocity and **(C)** movement offset for the three conditions. Dots represent individual participants. **(INSET)** Covariance plots of perpendicular deviations for a representative participant, overlaid across conditions at **(B)** peak velocity and **(C)** movement offset. **(D, E)** Covariance between perpendicular deviations of the right and left arms at **(D)** peak velocity and **(E)** movement offset for the three gain conditions. Each panel shows data from all trials across all participants. Darker ellipses denote 95% confidence ellipses of group-level data, and dark lines indicate group-level orthogonal regression fits (annotated values). Lighter ellipses reflect confidence ellipses for individual participants. Dashed line (slope = –45°) denotes the line of equivalence. Positive deviations in the left arm and negative deviations in the right arm represent medial perpendicular deviations **(F, G)** Bootstrapped distributions (10,000 resamples) of the overall, group-level orthogonal regression slopes for the two asymmetric gain conditions (LH Perp–RH Para High: purple; RH Perp–LH Para High: green), estimated at **(F)** peak velocity and **(G)** movement offset. Bias-corrected and accelerated (BCa) 95% confidence intervals are presented and annotated above each distribution. Non-overlapping intervals indicate statistically distinct coordination strategies between the two conditions.

As in our previous experiments, we fit orthogonal regression models to the hand deviations, and computed 95% confidence ellipses for both individual and group deviation data in each condition, separately for perpendicular and parallel directions. Figures 6B and 6C shows the individual regression slopes for perpendicular deviations at peak velocity and movement offset respectively, with insets depicting changes for a single participant. Consistent with experiment 1, we observed task-dependent changes in slopes in the asymmetric conditions compared to Equal_1, at both time points. This trend in individual slopes was statistically confirmed at peak velocity, where a linear mixed-effects model revealed a significant effect of gain (Fixed effect slope = –34.24°, 95% CI = [−56.39°, −12.08°], p = 0.005, Figure 6B). However, at movement offset, the model did not reveal significant differences between gain conditions, likely due to higher variability (Fixed effect slope = –14.93°, 95% CI = [−40.49°, 10.63°], p = 0.231, Figure 6C).

Figure 6D shows the group-level covariance relationship between the perpendicular deviations of the two arms, measured at peak velocity. Our analysis revealed distinct coordination patterns in the three conditions: Equal_1, LH Perp–RH Para High and RH Perp–LH Para High. In the Equal_1 condition, where both arms contributed equally to the perpendicular and parallel motion of the cursor, the regression slope was −40.78°, closely aligning with the line of equivalence. In the LH Perp–RH Para High condition, the regression slope shifted to −5.48°, indicating constrained perpendicular deviations of the left arm and more variable perpendicular deviations of the right arm. In the RH Perp–LH Para High condition, the slope changed to - 68.11°, indicating restricted lateral movements of the right arm when compared to the left arm, which moved more freely. These slopes were consistent with the slopes of perpendicular deviations observed in experiment 1. Notably, these task-sensitive coordination patterns in the perpendicular deviations of the two arms were also evident at movement offset with slope values being comparable to those observed at peak velocity (Figure 6E). The difference in group slope values for the two asymmetric gain conditions was confirmed using our bootstrapping analysis. At peak velocity (Figure 6F), the 95% confidence intervals for LH Perp– RH Para High ([–10.29°, –0.01°]) and RH Perp–LH Para High ([–80.95°, –59.35°]) showed no overlap. This was also true at movement offset (LH Perp–RH Para High CI = [−11.57°, −4.85°], RH Perp–RH Para High CI = [−75.26°, −61.98°], Figure 6G), confirming statistically significant differences in coordination patterns and thereby, in the control strategies governing movements of the two arms.

We next examined how asymmetrically altering the parallel contribution of each arm to the cursor influenced the coordination patterns along movement extent. We first analyzed the cursor errors along the *y*-axis across successive 10-trial cycles to determine differences across gain conditions (Figure 7A). A Kruskal-Wallis test revealed significant effects of gain in cycle 1 (p = 0.0002, Table 2); post-hoc tests indicated that the LH Perp–RH Para High (p = 0.0016) and RH Perp–LH Para High (p = 0.0006) conditions differed significantly from Equal_1, but not from each other (p = 1.00). We then assessed coordination patterns using orthogonal regression models on the parallel deviations. Figures 7B and 7C show regression slopes for individual participants at peak velocity and movement offset, respectively, with insets depicting slope changes in a representative participant. Task-dependent differences in coordination were evident across conditions at movement offset but not at peak velocity. A linear mixed model revealed no significant effect of gain condition on individual slope angles measured at peak velocity (Fixed effect slope = −2.24°, 95% CI = [−6.76°, 2.28°], p = 0.314, Figure 7B) but a marginal effect was present at movement offset (Fixed effect slope = −8.64°, 95% CI = [−17.33°, 0.06°], p = 0.051, Figure 7C).

**Figure 7:**
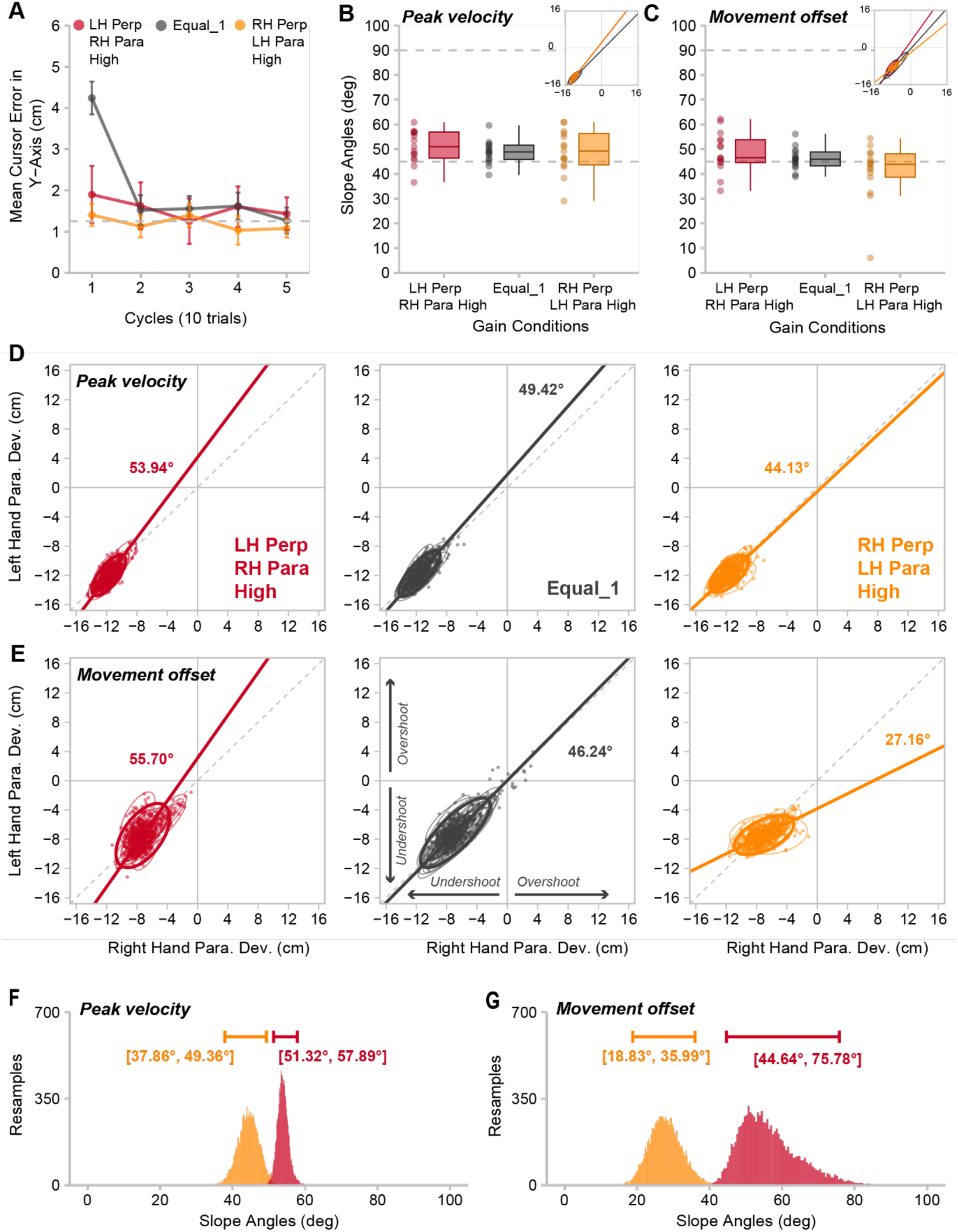
(A) Mean±SEM shared cursor error along the y-axis at movement offset, across five trial cycles, for the three gain conditions: LH Perp–RH Para High (red), Equal_1 (grey), and RH Perp–LH Para High (orange). Positive values indicate overshoot; negative values indicate undershoot of the shared cursor with respect to the center of the target. Dashed line at 1.25 cm indicates the upper vertical boundary of the target **(B, C)** Boxplots of regression slopes computed for the parallel deviations of the right and left arms at **(B)** peak velocity and **(C)** movement offset for the three conditions. Dots represent individual participants. **(INSET)** Covariance plots of parallel deviations for a representative participant, overlaid across conditions at **(B)** peak velocity and **(C)** movement offset. **(D, E)** Covariance between parallel deviations of the right and left arms at **(D)** peak velocity and **(E)** movement offset for the three gain conditions. Each panel shows data from all trials across all participants. Darker ellipses denote 95% confidence ellipses of group-level data, and the dark lines indicate group-level orthogonal regression fits (annotated values). Lighter ellipses reflect confidence ellipses for individual participants. Dashed line (slope = 45°) denotes the line of equivalence. Positive deviations indicate overshoot, and negative deviations indicate undershoot relative to the virtual target. **(F, G)** Bootstrapped distributions (10,000 resamples) of the overall, group-level orthogonal regression slopes for the two asymmetric gain conditions (LH Perp–RH Para High: red; RH Perp–LH Para High: orange), estimated at **(F)** peak velocity and **(G)** movement offset. Bias-corrected and accelerated (BCa) 95% confidence intervals are presented and annotated above each distribution. Non-overlapping intervals indicate statistically distinct coordination strategies between the two conditions.

Group-level covariance relationships between the parallel deviations of the two arms are shown in Figure 7D (peak velocity) and 7E (movement offset). At peak velocity, consistent with Experiment 2, task-dependent modulation was less prominent in the RH Perp–LH Para High condition, which showed a slope of 44.13°, comparable to the Equal_1 (49.42°) condition. In contrast, in the LH Perp–RH Para High condition, where the right arm had higher parallel contribution, changes in spatial covariance were noticeable at peak velocity itself (53.94°). At movement offset (Figure 7E), we observed that the modulation of regression slopes in the asymmetric conditions became more pronounced. In the Equal_1 condition, the slope was 46.24°. The two arms also consistently moved less, with mean movement extent of 8.87 cm in the left arm and 9.14 cm in the right arm to compensate for the increased cursor error in cycle 1. In the LH Perp–RH Para High condition, the regression slope for parallel deviations (55.70°) was not very different from that observed in RH Para High condition of experiment 2, indicating that the right arm had more constrained parallel variability than the left arm. In RH Perp–LH Para High condition, the slope (27.16°), resembled the LH Para High condition from experiment 2. This revealed more constrained parallel variability of the left arm while the right arm was allowed to be more variable. This trend at the group-level was statistically confirmed using bootstrapping analysis. The 95% confidence intervals of the LH Perp–RH Para High and RH Perp–LH Para High conditions showed no overlap at either peak velocity (LH Perp–RH Para High: [51.32°, 57.89°], RH Perp–LH Para High: [37.86°, 49.36°], Figure 7F) or movement offset (LH Perp–RH Para High: [44.64°, 75.78°], RH Perp–LH Para High: [18.83°, 35.99°], Figure 7G), indicating significant coordination differences between these conditions.

In sum, the changes in coordination in experiment 3 showed similarities to the patterns observed for perpendicular deviations in experiment 1 and parallel deviations in experiment 2. We also noted that participants demonstrated rapid adaptation to the combined perturbations within 20 trials for both perpendicular and parallel deviations in each condition (See exp. 3 Supplementary Results, Figures S3 and S4).

### Experiment 4

The previous experiment demonstrated that when perpendicular and parallel gains were applied simultaneously but asymmetrically to the arms, the arm with the higher perpendicular contribution exhibited reduced lateral variability, while the arm with the higher parallel contribution showed restricted parallel variability. In contrast, the arms with the lower contribution to extent and direction dimensions of the shared cursor moved more freely in those dimensions. However, it remained unclear whether this task-dependent coordination would hold when only one arm is primarily responsible for the control of both perpendicular and parallel motion of the cursor. We therefore proceeded to our fourth experiment, where we examined this issue by having one arm contribute more to both perpendicular and parallel cursor motion, while the other arm contributed less in both directions. In the LH Perp–Para High condition, the left arm had a higher contribution to both the perpendicular and parallel motion of the cursor, while the right arm had a lower contribution to both. Conversely, in the RH Perp–Para High condition, the right arm contributed more to the perpendicular and parallel motion of the cursor, while the left arm contributed less. We predicted that the arm with the higher contribution would show reduced variability in *both* dimensions.

We first analysed the mean endpoint cursor errors along the *x*-axis across 10-trial cycles to determine whether they differed across conditions (Figure 8A). A Kruskal-Wallis test revealed no significant differences across gain conditions in any cycle (all p > 0.10, Table 2). We separately examined the covariance for perpendicular and parallel deviations at the individual and group-level across conditions. Figures 8B and 8C show the individual regression slopes for perpendicular deviations at peak velocity and movement offset, respectively, with insets showing how the slopes changed across conditions in a representative participant. While some deviation in slopes from the Equal_1 condition was evident at peak velocity, the differences became a bit more pronounced at movement offset. However, across individuals, a linear mixed model revealed no significant effect of gain condition on slopes either at peak velocity (Fixed effect slope = –1.92°, 95% CI = [−25.02°, 21.19°], p = 0.861, Figure 8B) or at movement offset (Fixed effect slope = –2.40°, 95% CI = [−26.91°, 22.12°], p = 0.837, Figure 8C).

**Figure 8:**
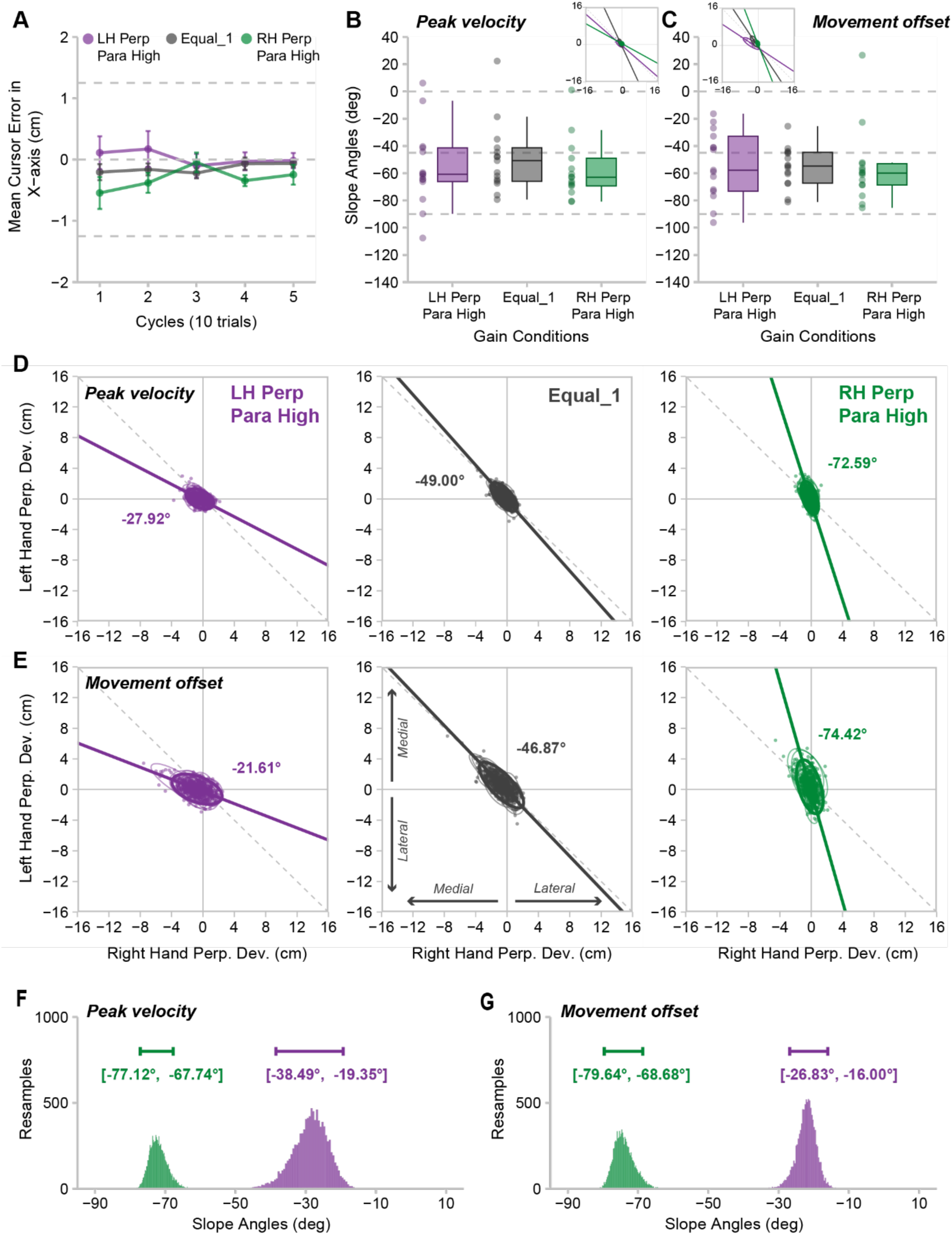
(A) Mean±SEM shared cursor error along the x-axis at movement offset across five cycles plotted for the three gain conditions: LH Perp–Para High (purple), Equal_1 (grey), and RH Perp–Para High (green). Positive values indicate rightward errors with respect to the center of the target; negative values indicate leftward errors. Dashed lines at ±1.25 cm mark the lateral boundaries of the target. **(B, C)** Boxplots of regression slopes computed for the perpendicular deviations of the right and left arms at **(B)** peak velocity and **(C)** movement offset for the three conditions. Dots represent individual participants. **(INSET)** Covariance plots of perpendicular deviations for a representative participant, overlaid across conditions at **(B)** peak velocity and **(C)** movement offset. **(D, E)** Covariance between perpendicular deviations of the right and left arms at **(D)** peak velocity and **(E)** movement offset for the three gain conditions. Each panel shows data from all trials across all participants. Darker ellipses denote 95% confidence ellipses of group-level data, and the dark lines indicate group-level orthogonal regression fits (annotated values). Lighter ellipses reflect confidence ellipses for individual participants. Dashed line (slope = –45°) denotes the line of equivalence. Positive deviations in the left arm and negative deviations in the right arm represent medial perpendicular deviations. **(F, G)** Bootstrapped distributions (10,000 resamples) of the overall, group-level orthogonal regression slopes for the two asymmetric gain conditions (LH Perp–Para High: purple; RH Perp–Para High: green), estimated from at **(F)** peak velocity and **(G)** movement offset. Bias-corrected and accelerated (BCa) 95% confidence intervals are presented and annotated above each distribution. Non-overlapping intervals indicate statistically distinct coordination strategies between the two conditions.

The group-level covariance relationships for the perpendicular deviation at peak velocity and movement offset are shown in Figures 8D and 8E, respectively. Group-level analysis revealed results consistent with experiment 3, demonstrating clear shifts in regression slopes in both gain conditions relative to the Equal_1 condition. As was the case before, these coordination patterns emerged early, and were already evident at peak velocity (Figure 8D). At movement offset, in the Equal_1 condition, perpendicular deviations exhibited negative covariation (slope = –46.87°, Figure 8E). However, in the LH Perp–Para High condition, regression slopes changed to –21.61°, implying that variance became constrained along the perpendicular axis for the left arm, while the right arm moved freely. On the contrary, in the RH Perp–Para High condition, the slope was –74.42, suggesting that it was the variance of the right arm that was now constrained, with the left arm being allowed to vary freely. We confirmed these observations with our bootstrapping analysis. For the perpendicular deviations, the confidence intervals between the LH Perp–Para High and RH Perp–Para High conditions did not overlap at peak velocity (LH Perp–Para High: CI = [–38.49°, –19.35°], RH Perp–Para High: [–77.12°, –67.74°], Figure 8F). This was also the case at movement offset (LH Perp–Para High: [–26.83°, –16.00°] and RH Perp–Para High: [–79.64°, –68.68°], Figure 8G), indicating robust task-dependent modulation of interlimb coordination.

To test for changes in coordination patterns along movement extent, we analyzed errors along the *y*-axis and changes in covariance relationships across conditions. Analysis of mean endpoint cursor errors along the *y*-axis (Figure 9A) revealed a significant effect only in cycle 1 (p < 0.0001); post-hoc comparisons revealed that both LH Perp–Para High (p < 0.0001) and RH Perp–Para High (p < 0.0001) differed significantly from Equal_1, but not from each other (p = 1.00). Figures 9B and 9C show the individual regression slopes for parallel deviations at peak velocity and movement offset respectively, with insets depicting slope changes in one participant. Task-dependent differences in coordination were not prominent at peak velocity but were more apparent at movement offset. We used a linear mixed-effects model to confirm this trend observed in individual slopes. Only marginal effects of gain condition on slope angles at peak velocity (Fixed effect slope = 3.83°, 95% CI = −0.18°, 7.83°], *p* = 0.060, Figure 9B) and at movement offset (Fixed effect slope = 11.53°, 95% CI = [−0.78°, 23.84°], p = 0.064, Figure 9C) were observed.

**Figure 9:**
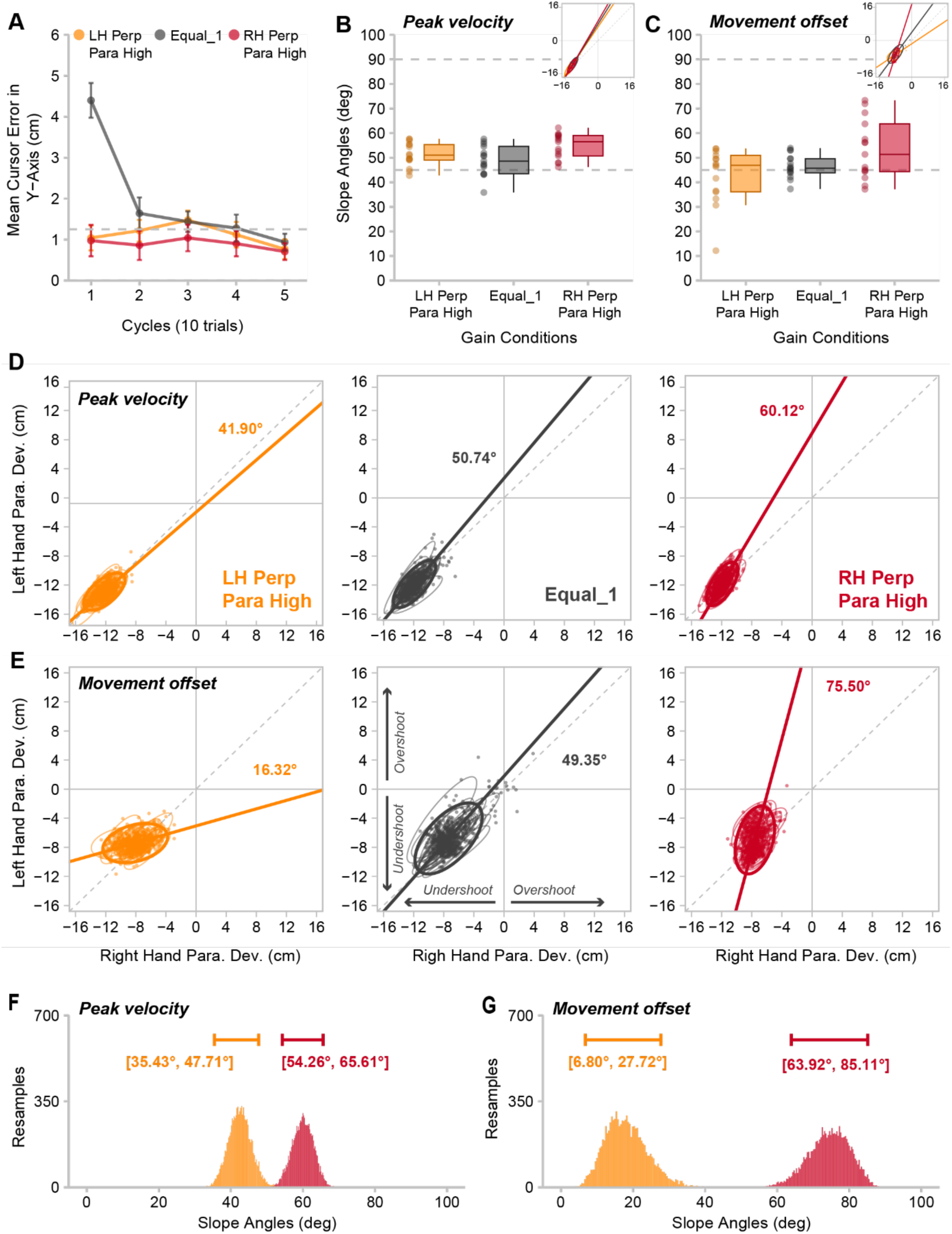
(A) Mean±SEM shared cursor error along the y-axis at movement offset, across five trial cycles, for the three gain conditions: LH Perp–Para High (orange), Equal_1 (grey), and RH Perp–Para High (red). Positive values indicate overshoot; negative values indicate undershoot of the shared cursor with respect to the target center. Dashed line at 1.25 cm indicates upper vertical boundary of the target. **(B, C)** Boxplots of regression slopes computed for the parallel deviations of the right and left arms at **(B)** peak velocity and **(C)** movement offset for the three conditions. Dots represent individual participants. **INSET** Covariance plots of parallel deviations for a representative participant, overlaid across conditions at **(B)** peak velocity and **(C)** movement offset. **(D, E)** Covariance between parallel deviations of the right and left arms at **(D)** peak velocity and **(E)** movement offset for the three gain conditions. Each panel shows data from all trials across all participants. Darker ellipses denote 95% confidence ellipses of group-level data, and the dark lines indicate group-level orthogonal regression fits (annotated values). Lighter ellipses reflect confidence ellipses for individual participants. Dashed line (slope = 45°) denotes the line of equivalence. Positive deviations indicate overshoot, and negative deviations indicate undershoot relative to the virtual target. **(F, G)** Bootstrapped distributions (10,000 resamples) of the overall, group-level orthogonal regression slopes for the two asymmetric gain conditions (LH Perp–Para High: orange; RH Perp–Para High: red), estimated at **(F)** peak velocity and **(G)** movement offset. Bias-corrected and accelerated (BCa) 95% confidence intervals are presented and annotated above each distribution. Non-overlapping intervals indicate statistically distinct coordination strategies between the two conditions.

Analysis of group-level covariance relationships between the parallel deviations of the left and right arm at peak velocity and movement offset are shown in Figures 9D and 9E respectively. Changes in coordination at peak velocity were observed only when the right arm had higher contribution to the parallel motion of the shared cursor (RH Perp–Para High slope = 60.12°), but not in the LH Perp–Para High (41.90°) condition. However, these changes were evident in both conditions at movement offset. In the LH Perp–Para High condition, the regression slope for parallel deviations changed to 16.32°, reflecting constrained parallel variability in the left arm. In the RH Perp–Para High condition, the slope was modulated to 75.50°, implying restricted parallel variability in the right arm. As before, since individual slopes were not reliable, we bootstrapped the group-level slopes to statistically assess differences in coordination patterns at peak velocity and movement offset. This analysis revealed no overlap of confidence intervals between LH Perp–Para High ([35.43°, 47.71°]) and RH Perp–Para High ([54.26°, 65.61°]) conditions at peak velocity (Figure 9F). Similarly, no overlap of confidence intervals was observed at movement offset either (LH Perp–Para High: [6.80°, 27.72°] and RH Perp– Para High: [63.92°, 85.11°], Figure 9G), suggesting clearly different coordination strategies in these two conditions.

Finally, to examine the timescale of adaptation, orthogonal regression models were fit to successive 10-trial cycles for both the asymmetric gain conditions. Consistent with previous experiments, rapid adaptation was observed in the covariance relationships, within 10 – 20 trials of introducing the asymmetric gains (See exp. 4 Supplementary Results, Figures S5 and S6).

## DISCUSSION

Effective coordination between the upper limbs is essential to perform skilled bimanual actions, but the control policy underlying such coordination remains debated. Early research (Kelso et al., 1979a; 1979b) suggested rigid coupling of homologous muscle groups for bimanual coordination, but more recent work has pointed to a more flexible control framework, shaped by the goal and demands of the task (Diedrichsen, 2007; Diedrichsen and Gush, 2009; Mutha and Sainburg, 2009; Ranganathan et al., 2019; Kitchen et al. 2023; Yuk et al., 2024). In this study, we demonstrate that this task-dependent flexibility extends to distinct movement axes, particularly movement direction and extent, implying that the sensorimotor system is capable of adjusting coordination largely independently along these dimensions based on task demands. This adaptive behaviour was evident as a change in movement variability at peak velocity in some cases but most certainly at movement end. Specifically, when one arm contributed more to the perpendicular motion of the cursor, its variability in that direction was reduced compared to the arm that contributed less. Likewise, when we altered each arm’s contribution to the parallel motion of the cursor, variability along the parallel axis was reduced in the arm that contributed more while the other arm exhibited greater variability. Furthermore, when we simultaneously altered the contribution of each arm to both perpendicular and parallel motion, we again found that the arm with higher perpendicular contribution to the cursor consistently showed reduced lateral variability, while the arm with the higher parallel contribution demonstrated restricted parallel variability. The corresponding lower contribution arms exhibited higher variability, suggesting a compensatory role in stabilizing the shared cursor. Taken together, our findings suggest that the sensorimotor system is able to resolve the shared cursor errors across movement direction and extent, and flexibly coordinate the two arms specific to the demands along that corresponding axis.

We arrived at this conclusion primarily through our group-level analysis of regression slopes. We observed high variability in individual slope estimates for both perpendicular and parallel deviations across all our experiments. This could, in part, have stemmed from limitations associated with fitting regression model to a smaller number of data points. The individual slopes were estimated using orthogonal regression model fit to only 50 data points per condition. While orthogonal regression accounts for measurement errors in both variables used in the fit, its root mean squared error (RMSE) is known to increase when the number of data points is smaller (Tod et al., 2002), resulting in a poorer fit. This made our individual slope estimates more susceptible to the influence of outliers and less reliable overall. Given this limitation, we considered our group-level slopes, computed using orthogonal regression fits to a substantially larger set of 750 data points per condition, to be a more robust and reliable measure. To further ensure the reliability of these group-level findings and to compare the flexible changes in coordination between conditions, we also bootstrapped the group-level slopes. This approach helped us confirm that our conclusions were based on stable, well-supported data, mitigating the limitations of individual-level analyses.

We observed an intriguing interlimb difference in the spatial covariance of parallel deviations when we altered the contribution of each arm to the parallel motion of the cursor either independently (in experiment 2) or simultaneously (in experiments 3 and 4). Specifically, when the left arm had a higher parallel contribution (LH Para High, RH Perp–LH Para High and LH Perp–Para High conditions), the covariance between the parallel deviations at peak velocity was comparable to that of the Equal_1 condition. However, when the right arm had a higher parallel contribution (RH Para High, LH Perp–RH Para High and RH Perp–Para High conditions), the covariance of the parallel deviations at peak velocity deviated from Equal_1 and resembled their corresponding pattern observed at movement offset. These differences could be related to the differences in control mechanisms that each arm-hemisphere system is thought to be specialized for (Woytowicz et al., 2018); it has been shown in unimanual reaching in right-handers that movements of the right (dominant) arm are driven largely through predictive (feedforward) control mechanisms, while movements of the left (non-dominant) arm are more reliant on feedback processes (Sainburg and Schaefer, 2004; Sainburg, 2005; Przybyla et al., 2011; Mutha et al., 2013). It is therefore possible that when the right arm has a higher contribution along movement extent, the sensorimotor system engages predominantly feedforward control mechanisms to pre-emptively correct for errors, resulting in coordination changes that can be seen early on. In contrast, when the left arm has a higher contribution to cursor motion, the adjustments are likely driven by online feedback, and therefore take longer to emerge. In line with this idea, Schaffer et al. (2020) have shown that right-handed individuals with left-hemisphere damage exhibit impaired bilateral coordination before peak velocity during a bimanual bar-moving task, while right-hemisphere damage does not disrupt this predictive phase, suggesting that feedforward coordination of both limbs relies heavily on intact left-hemisphere mechanisms, possibly influencing control of the right arm more.

This lateralized response at peak velocity was, however, not observed in the perpendicular deviations when we altered perpendicular gain. The slopes of the perpendicular deviations in the two asymmetric conditions at peak velocity were already close to the slopes observed at movement offset, suggesting that predictive mechanisms might have been engaged even when the left arm had a greater contribution to the cursor’s perpendicular motion. One possible explanation for this could be that the sensorimotor system may have more effectively updated its internal models using the signed cursor errors along the perpendicular dimension (*x*-axis). Assuming that perpendicular corrections are primarily driven by these directional errors, the two asymmetric conditions, which produce errors in opposite directions, would provide clearer, direction-specific feedback. This then likely enables the system to incorporate feedback better to generate more precise predictions about the required movement direction for each arm, facilitating effective feedforward adjustments. As a result, lateralized differences in control may have been minimized during the early movement phase. In contrast, cursor errors along the *y*-axis were consistently positive (overshooting) in the two asymmetric gain cases, thereby providing minimal information about the directionality of errors. This may also explain why the differences in perpendicular deviations between the asymmetric conditions at movement offset were more pronounced than those seen for parallel deviations.

It is interesting to note that adaptive changes were seen following perpendicular gain manipulations in Experiments 1, 3 and 4 even though the shared cursor, on average, fell within the lateral boundaries of the target leading to task success. However, we observed that considerable trial-to-trial variability was present and the cursor did not always hit the target, which may have provided necessary error signals for adaptive changes in coordination. Despite mean success, participants frequently missed the target center, generating directional endpoint errors that deviated leftward or rightward depending on the asymmetric gain condition. These errors may have been used to continuously update internal models based on reach errors (Diedrichsen et al., 2005, Krakauer et al., 2000, Wolpert et al., 2011). Indeed, prior work suggests that the presence of endpoint variability offers more informative error signals than mean accuracy alone (Wu et al., 2014; Dhawale et al., 2017; Sternad, 2018), and that inconsistent target hits can elicit learning on the next trial. Thus, even when average performance appeared successful, trial-wise errors likely played a critical role in driving the observed flexible adjustments in coordination, particularly in the perpendicular direction.

More broadly, our findings provide compelling evidence for independent yet interacting control mechanisms governing movement extent and direction during bimanual coordination. The clearest demonstration of independence emerges from experiments 1 and 2, where altering arm contributions along only one dimension produced no changes in the unaltered dimension, indicating that the sensorimotor system can selectively modulate coordination strategies without cross-dimensional interference. This axis-specific control extended to experiments 3 and 4, where simultaneous alterations of both direction and extent gains revealed that the system coordinated the two arms flexibly and independently along each axis according to task constraints. The independence of extent and direction control is further supported by lateralized control strategies observed in parallel (extent) deviations, but not in perpendicular (direction) deviations, where the timing of these modulations appeared early in the movement along extent in right hand high conditions, but only at movement completion for left hand high conditions.

However, our data also point to some interaction between extent and direction control, rather than complete independence. In experiments 3 and 4, where both movement dimensions were manipulated simultaneously, we observed greater variability in the individual regression slopes of perpendicular deviations across participants compared to experiment 1, where only direction was manipulated. This increased variability may reflect the added complexity of simultaneously controlling extent, which could have interfered with the effective use of visual feedback for directional error correction (Sarlegna & Blouin, 2010). Specifically, in experiment 3, decreasing the extent contribution of the arm with higher direction contribution reversed the sign of cursor errors along the *x*-axis between the two asymmetric conditions compared to those in experiment 1. This was not observed in experiment 4, when the same arm had high perpendicular and parallel contribution to the cursor. Moreover, cursor errors along the *x*-axis decreased in experiment 3 and 4, compared to experiment 1, where only directional gains were applied. All of these observations suggest that extent constraints influenced how the system coordinated movement direction. Furthermore, the absolute difference at movement offset in perpendicular deviation between the asymmetric conditions was greater in experiment 3 (61.41) than in experiment 4 (52.81), while parallel deviations showed the opposite pattern (28.54 versus 59.18 respectively). This suggests that applying both gains to the same arm in experiment 4 enhanced extent coordination at the cost of directional precision, consistent with evidence that amplitude control is more susceptible to directional interference than vice versa (Pan and Van Gemmert, 2019). Taken together, our findings reinforce the idea that extent and direction rely on partially distinct yet interacting control pathways (Wenderoth et al., 2005; Wijdenes et al., 2013), and that coordinating both dimensions simultaneously can modulate the effectiveness of task-specific strategies.

Our results also offer insight into the control policy underlying bimanual coordination. We made two primary observations: 1) task-specific changes in coordination when the contributions of each arm to the shared cursor were asymmetric, and 2) reduced variability in the arm with higher contribution to the cursor, with a corresponding increase in variability in the lower-contribution arm. The first observation challenges models that assume a rigid, bilaterally coupled control mechanism. If the coupling hypothesis were strictly true, then we would expect that spatial symmetry between the limbs would be preserved even when peripheral conditions changed. Instead, we found that participants flexibly adjusted coordination strategies to match task demands, modulating variability asymmetrically based on the contribution of each arm to the shared feedback cursor. The second observation aligns with the “minimum intervention principle” of optimal feedback control theory, which posits that the motor system allows variability to accumulate in task-irrelevant dimensions while actively constraining variability in those dimensions that are relevant for task success. (Todorov and Jordan, 2002; Scott, 2004; Diedrichsen et al., 2010). In our study, the arm with lower contribution represents the “task-irrelevant” dimension, while the arm with higher contribution constitutes the “task-relevant” dimension that directly impacts cursor performance. In line with the optimal feedback control hypothesis, the system prioritized reducing variability in the specific dimension and specific arm that had a higher contribution to the task. These results suggest that bimanual coordination is governed by flexible task-dependent control policies, wherein the system adapts in real-time based on the relative task relevance of each limb along each movement axis.

Finally, it is important to point out that the covariance between the two arms remained non-zero under the asymmetric conditions. This suggests that coordination is not completely decoupled. The two arms responded to the applied visual feedback gain independently, thereby breaking the inherent co-dependency observed in shared cursor tasks (Diedrichsen, 2007; Mutha and Sainburg, 2009). However, these adjustments were not entirely independent: reductions in variability in one arm were accompanied by compensatory increases in the other. This pattern of correlated changes in the two arms and the residual covariation indicate that some level of shared, coordinated control is maintained even when task demands favour asymmetry (Kitchen et al., 2023). Future studies could test the limits of this shared control through further modifications in task demands including changes in feedback, preparation time and practice.

## Supporting information

Supplementary Results

## Acknowledgements

We thank Ajay Kumar Sahu for assistance with the development of the data collection code. We are also thankful to Dr. Nick Kitchen, Dr. Robert Sainburg, Gaurav Panthi and Sinchana Vaasanthi for many helpful discussions regarding this study. This work was supported by grants from the Department of Science and Technology, Government of India. Support from IIT Gandhinagar is also acknowledged.

## Notes

**Conflicts of interest:** The authors declare no competing financial interests.

### Competing Interest Statement

The authors have declared no competing interest.

## REFERENCES

D’Cruz, N., De Vleeschhauwer, J., Putzolu, M., Nackaerts, E., Gilat, M., & Nieuwboer, A. (2024). Sensorimotor network segregation predicts Long-Term learning of writing skills in Parkinson’s disease. Brain Sciences, 14(4), 376. 10.3390/brainsci14040376

Dhawale, A. K., Smith, M. A., & Ölveczky, B. P. (2017). The role of variability in motor learning. Annual Review of Neuroscience, 40(1), 479–498. 10.1146/annurev-neuro-072116-031548

Diedrichsen, J. (2007). Optimal Task-Dependent changes of bimanual feedback control and adaptation. Current Biology, 17(19), 1675–1679. 10.1016/j.cub.2007.08.051

Diedrichsen, J., & Gush, S. (2009). Reversal of bimanual feedback responses with changes in task goal. Journal of Neurophysiology, 101(1), 283–288. 10.1152/jn.90887.2008

Diedrichsen, J., Hashambhoy, Y., Rane, T., & Shadmehr, R. (2005). Neural correlates of reach errors. Journal of Neuroscience, 25(43), 9919–9931. 10.1523/jneurosci.1874-05.2005

Diedrichsen, J., Shadmehr, R., & Ivry, R. B. (2010). The coordination of movement: optimal feedback control and beyond. Trends in Cognitive Sciences, 14(1), 31–39. 10.1016/j.tics.2009.11.004

Dimitriou, M., Franklin, D. W., & Wolpert, D. M. (2012). Task-dependent coordination of rapid bimanual motor responses. Journal of Neurophysiology, 107(3), 890–901. 10.1152/jn.00787.2011

Favilla, M., Gordon, J., Hening, W., & Ghez, C. (1990). Trajectory control in targeted force impulses: VII. Independent setting of amplitude and direction in response preparation. Experimental Brain Research, 79(3), 530–538. 10.1007/bf00229322

Favilla, M., Hening, W., & Ghez, C. (1989). Trajectory control in targeted force impulses: VI. Independent specification of response amplitude and direction. Experimental Brain Research, 75(2), 280–294. 10.1007/bf00247934

Franz, E. A. (1997). Spatial coupling in the coordination of complex actions. The Quarterly Journal of Experimental Psychology Section A, 50(3), 684–704. 10.1080/713755726

Franz, E. A., Zelaznik, H. N., & McCabe, G. (1991). Spatial topological constraints in a bimanual task. Acta Psychologica, 77(2), 137–151. 10.1016/0001-6918(91)90028-X

Fu, Q. G., Flament, D., Coltz, J. D., & Ebner, T. J. (1995). Temporal encoding of movement kinematics in the discharge of primate primary motor and premotor neurons. Journal of Neurophysiology, 73(2), 836–854. 10.1152/jn.1995.73.2.836

Fu, Q. G., Suarez, J. I., & Ebner, T. J. (1993). Neuronal specification of direction and distance during reaching movements in the superior precentral premotor area and primary motor cortex of monkeys. Journal of Neurophysiology, 70(5), 2097–2116. 10.1152/jn.1993.70.5.2097

Gordon, J., Ghilardi, M. F., & Ghez, C. (1994). Accuracy of planar reaching movements: I. Independence of direction and extent variability. Experimental Brain Research, 99(1), 97–111. 10.1007/bf00241415

Hardwick, R. M., Rajan, V. A., Bastian, A. J., Krakauer, J. W., & Celnik, P. A. (2017). Motor learning in stroke. Neurorehabilitation and Neural Repair, 31(2), 178–189. 10.1177/1545968316675432

Kelso, J. S., Putnam, C. A., & Goodman, D. (1983). On the Space-Time Structure of Human Interlimb Co-Ordination. The Quarterly Journal of Experimental Psychology Section A, 35(2), 347–375. 10.1080/14640748308402139

Kelso, J. S., Southard, D. L., & Goodman, D. (1979a). On the coordination of two-handed movements. Journal of Experimental Psychology Human Perception & Performance, 5(2), 229–238. 10.1037/0096-1523.5.2.229

Kelso, J. a. S., Southard, D. L., & Goodman, D. (1979b). On the Nature of Human Interlimb Coordination. Science, 203(4384), 1029–1031. 10.1126/science.424729

Kitchen, N. M., Yuk, J., Przybyla, A., Scheidt, R. A., & Sainburg, R. L. (2023). Bilateral arm movements are coordinated via task-dependent negotiations between independent and codependent control, but not by a “coupling” control policy. Journal of Neurophysiology, 130(3), 497–515. 10.1152/jn.00501.2022

Krakauer, J. W., Pine, Z. M., Ghilardi, M., & Ghez, C. (2000). Learning of visuomotor transformations for vectorial planning of reaching trajectories. Journal of Neuroscience, 20(23), 8916–8924. 10.1523/JNEUROSCI.20-23-08916.2000

Marteniuk, R. G., MacKenzie, C. L., & Baba, D. M. (1984). Bimanual Movement Control: Information processing and Interaction Effects. The Quarterly Journal of Experimental Psychology Section A, 36(2), 335–365. 10.1080/14640748408402163

Messier, J., & Kalaska, J. F. (1997). Differential effect of task conditions on errors of direction and extent of reaching movements. Experimental Brain Research, 115(3), 469–478. 10.1007/pl00005716

Mutha, P. K., Boulinguez, P., & Sainburg, R. L. (2008). Visual modulation of proprioceptive reflexes during movement. Brain Research, 1246, 54–69. 10.1016/j.brainres.2008.09.061

Mutha, P. K., Haaland, K. Y., & Sainburg, R. L. (2013). Rethinking motor lateralization: specialized but complementary mechanisms for motor control of each arm. PLoS ONE, 8(3), e58582. 10.1371/journal.pone.0058582

Mutha, P. K., & Sainburg, R. L. (2007). Control of velocity and position in single joint movements. Human Movement Science, 26(6), 808–823. 10.1016/j.humov.2007.06.001

Mutha, P. K., & Sainburg, R. L. (2009). Shared bimanual tasks elicit bimanual reflexes during movement. Journal of Neurophysiology, 102(6), 3142–3155. 10.1152/jn.91335.2008

Omrani, M., Diedrichsen, J., & Scott, S. H. (2013). Rapid feedback corrections during a bimanual postural task. Journal of Neurophysiology, 109(1), 147–161. 10.1152/jn.00669.2011

Pan, Z., & Van Gemmert, A. W. (2019). The control of amplitude and direction in a bimanual coordination task. Human Movement Science, 65, 111–120. 10.1016/j.humov.2018.03.014

Przybyla, A., Haaland, K. Y., Bagesteiro, L. B., & Sainburg, R. L. (2011). Motor asymmetry reduction in older adults. Neuroscience Letters, 489(2), 99–104. 10.1016/j.neulet.2010.11.074

Puth, M., Neuhäuser, M., & Ruxton, G. D. (2015). On the variety of methods for calculating confidence intervals by bootstrapping. Journal of Animal Ecology, 84(4), 892–897. 10.1111/1365-2656.12382

Ranganathan, R., Gebara, R., Andary, M., & Sylvain, J. (2019). Chronic stroke survivors show task-dependent modulation of motor variability during bimanual coordination. Journal of Neurophysiology, 121(3), 756–763. 10.1152/jn.00218.2018

Sainburg, R. L. (2005). Handedness: differential specializations for control of trajectory and position. Exercise and Sport Sciences Reviews, 33(4), 206–213. 10.1097/00003677-200510000-00010

Sainburg, R. L., & Schaefer, S. Y. (2004). Interlimb differences in control of movement extent. Journal of Neurophysiology, 92(3), 1374–1383. 10.1152/jn.00181.2004

Sarlegna, F. R., & Blouin, J. (2010). Visual guidance of arm reaching: Online adjustments of movement direction are impaired by amplitude control. Journal of Vision, 10(5), 24. 10.1167/10.5.24

Schaffer, J. E., Maenza, C., Good, D. C., Przybyla, A., & Sainburg, R. L. (2020). Left hemisphere damage produces deficits in predictive control of bilateral coordination. Experimental Brain Research, 238(12), 2733–2744. 10.1007/s00221-020-05928-2

Scott, S. H. (2004). Optimal feedback control and the neural basis of volitional motor control. Nature Reviews. Neuroscience, 5(7), 532–545. 10.1038/nrn1427

Shingane, S. N., Rao, N., Kumar, N., & Mutha, P. K. (2025). Task relevance selectively modulates sensorimotor adaptation in the presence of multiple prediction errors. Journal of Neurophysiology. 10.1152/jn.00511.2024

Sternad, D. (2018). It’s not (only) the mean that matters: variability, noise and exploration in skill learning. Current Opinion in Behavioral Sciences, 20, 183–195. 10.1016/j.cobeha.2018.01.004

Tod, M., Aouimer, A., & Petitjean, O. (2002). Estimation of pharmacokinetic parameters by orthogonal regression: comparison of four algorithms. Computer Methods and Programs in Biomedicine, 67(1), 13–26. 10.1016/S0169-2607(00)00148-6

Todorov, E., & Jordan, M. I. (2002). Optimal feedback control as a theory of motor coordination. Nature Neuroscience, 5(11), 1226–1235. 10.1038/nn963

Wenderoth, N., Debaere, F., Sunaert, S., & Swinnen, S. P. (2005). Spatial interference during bimanual coordination: Differential brain networks associated with control of movement amplitude and direction. Human Brain Mapping, 26(4), 286–300. 10.1002/hbm.20151

Wijdenes, L. O., Brenner, E., & Smeets, J. B. J. (2013). Comparing online adjustments to distance and direction in fast pointing movements. Journal of Motor Behavior, 45(5), 395–404. 10.1080/00222895.2013.815150

Wolpert, D. M., Diedrichsen, J., & Flanagan, J. R. (2011). Principles of sensorimotor learning. Nature Reviews. Neuroscience, 12(12), 739–751. 10.1038/nrn3112

Woytowicz, E. J., Westlake, K. P., Whitall, J., & Sainburg, R. L. (2018). Handedness results from complementary hemispheric dominance, not global hemispheric dominance: evidence from mechanically coupled bilateral movements. Journal of Neurophysiology, 120(2), 729–740. 10.1152/jn.00878.2017

Wu, H. G., Miyamoto, Y. R., Castro, L. N. G., Ölveczky, B. P., & Smith, M. A. (2014). Temporal structure of motor variability is dynamically regulated and predicts motor learning ability. Nature Neuroscience, 17(2), 312–321. 10.1038/nn.3616

Yuk, J., Kitchen, N. M., Przybyla, A., Scheidt, R. A., & Sainburg, R. L. (2024). Symmetry and synchrony of bimanual movements are not predicated on interlimb control coupling. Journal of Neurophysiology, 131(6), 982–996. 10.1152/jn.00476.2023

