## Supplementary Results for "Flexible, task-dependent bimanual coordination along movement direction and extent"

### Experiment 1

To probe the timescale over which participants adjusted to the altered hand-cursor relationships in the two asymmetric blocks, we analyzed how the covariance between the perpendicular deviations of the arms evolved over time at peak velocity and at movement offset. This was done by grouping the data into cycles of 10 successive trials and calculating the group covariance for each cycle. The slope angles for each cycle separately and the entire block cumulatively are provided in Table S1 and shown in Figure S1. At peak velocity (Figure S1-A), the group-level regression slopes in the first 10-trial cycle for the LH Perp High and RH Perp High conditions were -23.59° and -68.82°, respectively. This divergence from the Equal_1 slopes within the first few trials indicated rapid initial adjustment in coordination strategy upon the introduction of asymmetric perpendicular gains. By the third cycle in the LH Perp High (-15.09) and RH Perp High (-74.26°) slopes closely aligned with the overall group slopes calculated across the whole block (dashed purple and green lines, Figure S1-A), indicating that the rapid initial adaptation was stable and gradual throughout the block. Interestingly, by the final bin, the slopes further shifted to -10.46° and -77.41°, respectively, surpassing the overall group slope, suggesting continued adaptation to the imposed conditions. A similar trend was observed at movement offset (Figure S1-B), where changes in slope emerged within the first 10 trials and continued to evolve over the block. We did not conduct a timescale-based covariance analysis on the parallel deviations as gains along that axis were maintained at 1:1, and no meaningful change in group-level slopes was observed in the asymmetric conditions relative to the Equal_1 condition, at either peak velocity (Figure 3D) or at movement offset (Figure 3E).

***Table S1:*** *Regression slopes for each 10-trial cycle and entire block of perpendicular deviations between the two arms in Experiment 1*

| | At Peak Velocity | | At Movement Offset | |
|---|---|---|---|---|
| **Cycle** | **LH Perp High** | **RH Perp High** | **LH Perp High** | **RH Perp High** |
| **1 (1 - 10 trials)** | -23.59° | -68.82° | -21.97° | -70.11° |
| **2 (11 - 20 trials)** | -25.26° | -79.99° | -19.61° | -82.74° |
| **3 (21 - 30 trials)** | -15.09° | -74.26° | -13.74° | -73.16° |
| **4 (31 - 40 trials)** | -14.51° | -77.49° | -11.45° | -76.89° |
| **5 (41 - 50 trials)** | -10.46° | -77.41° | -5.83° | -77.74° |
| **Entire block** | -17.69° | -75.91° | -14.18° | -76.34° |

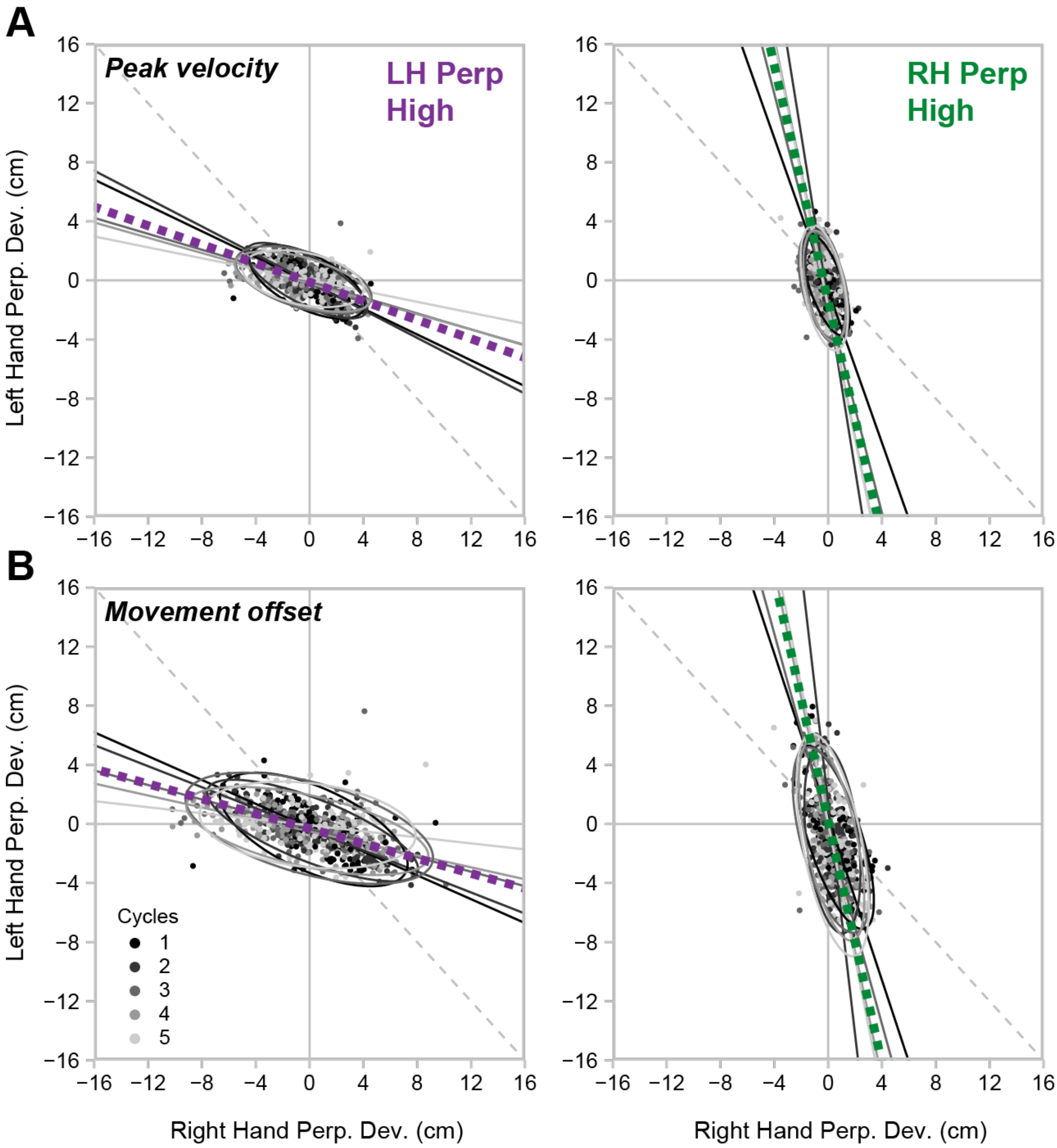


***Figure S1:*** *Group-level covariance relationships between the perpendicular deviations of the right and left arms over time in experiment 1, shown separately for LH Perp High (left column) and RH Perp High (right column) conditions at **(A)** peak velocity and **(B)** movement offset. Each subplot shows orthogonal regression lines and 95% confidence ellipses for successive 10-trial cycles, with shading from dark to light grey indicating progression across the block. Coloured dotted lines represent the overall regression line fit for the entire block (50 trials) in each condition, identical to the group-level slopes in Figure 2D and 2E. Dashed grey line (slope = –45°) denotes the line of equivalence.*

**Experiment 2**

We examined the timescale of adjustments in coordination to the asymmetric parallel gains by computing group-level slopes for the parallel deviations of the two arms across successive 10-trial cycles. The regression slopes for each cycle separately and entire block cumulatively, at peak velocity and at movement offset, are shown in Table S2. At peak velocity (Figure S2-A), the group-level slopes in all cycles, except cycle 3 (38.05°) and 4 (38.20°), oscillated around the line of equivalence in the LH Para High Condition, suggesting minimal changes in coordination early during the movement when the left arm had a higher contribution to the parallel motion of the cursor. In contrast, a clear shift in slope away from the Equal_1 slope was observed within the first 10 trials in the RH Para High condition, indicating task-dependent changes in coordination early in the movement when the right arm had higher contribution to the parallel motion of the cursor, consistent with the overall group-level slope (Figure 4D – RH Para High). At movement offset (Figure S2-B), we observed that participants in the LH Para High condition adapted rapidly, as evidenced by the group slope in the first 20 trials (30.14°) closely aligning with the overall group slope (31.67°). However, in the RH Para High condition, this adjustment took about 30 trials (58.49°) to align reasonably closely with the overall group-level slope (58.07°). Interestingly, the adaptation appeared to plateau in the LH Para High condition after 30 trials, whereas in the RH Para High condition participants continued to adapt throughout the block. This was reflected by the group-level slope in the last 10 trial-cycle (69.08°) surpassing the overall group-level slope, indicating continued adaptation. We did not perform this timescale analysis for perpendicular deviations of the two arms, as the perpendicular gain ratios were kept constant at 1:1 across conditions, and no significant differences in group-level slopes were observed between the asymmetric and Equal_1 conditions at either peak velocity (Figure 5D) or at movement offset (Figure 5E).

***Table S2:*** *Regression slopes for each 10-trial cycle and entire block of parallel deviations between the two arms in Experiment 2*

| | At Peak Velocity | | At Movement Offset | |
|---|---|---|---|---|
| **Cycle** | **LH Para High** | **RH Para High** | **LH Para High** | **RH Para High** |
| **1 (1 – 10 trials)** | 45.09° | 53.57° | 35.50° | 52.79° |
| **2 (11 – 20 trials)** | 42.63° | 54.79° | 30.14° | 54.47° |
| **3 (21 – 30 trials)** | 38.05° | 56.02° | 25.93° | 58.49° |
| **4 (31 – 40 trials)** | 38.20° | 56.31° | 32.05° | 60.47° |
| **5 (41 – 50 trials)** | 47.50° | 62.82° | 35.01° | 69.08° |
| **Entire Block** | 41.92° | 56.45° | 31.67° | 58.07° |

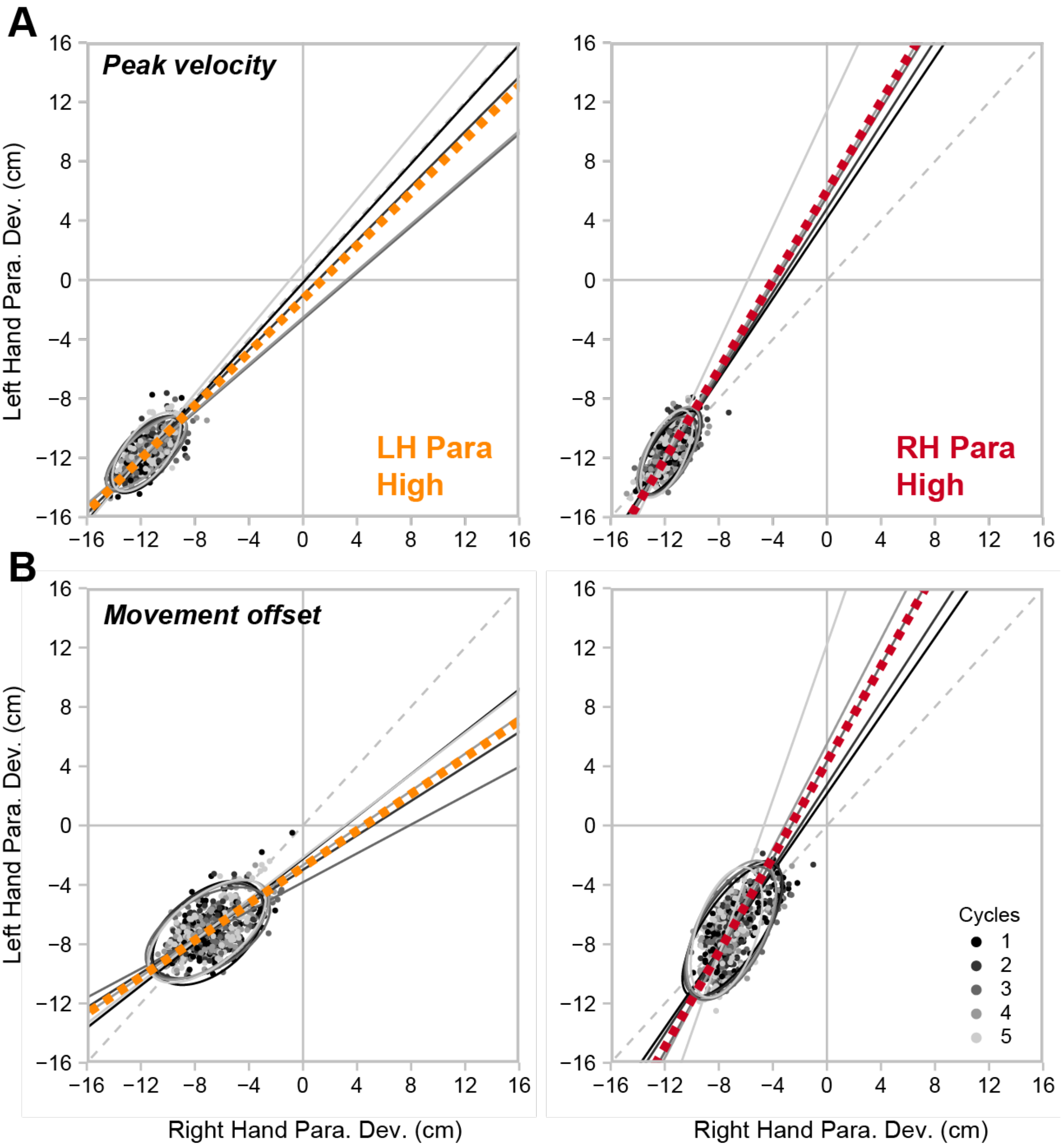


***Figure S2:*** *Group-level covariance relationships between the parallel deviations of the right and left arms over time in Experiment 2, shown separately for LH Para High (left column) and RH Para High (right column) conditions at* ***(A)*** *peak velocity and* ***(B)*** *movement offset. Each subplot shows orthogonal regression lines and 95% confidence ellipses for successive 10-trial cycles, with shading from dark to light grey indicating progression across the block. Coloured dotted lines represent the overall regression line fit for the entire block (50 trials) in each condition, identical to the group-level slopes in Figure 4D and 4E. Dashed grey line (slope = 45°) denotes the line of equivalence.*

**Experiment 3**

In this experiment, we simultaneously altered the contribution of each arm to both parallel and perpendicular motion of the shared cursor and observed task-dependent deviation of regression slopes in the two asymmetric LH Perp–RH Para High and RH Perp–LH Para High conditions relative to the Equal_1 condition in both the perpendicular deviations and parallel dimensions. The timescale over which these changes emerged in the asymmetric conditions were analyzed separately for the perpendicular and parallel deviations of the right and left arms. Table S3 shows the group-level slopes for each 10-trial cycle and entire block between perpendicular deviations of the two arms at peak velocity and at movement offset. At peak velocity (Figure S3-A), deviations from the Equal_1 slope emerged early in both conditions. By the second 10-trial cycle, group-level slopes in the LH Perp–RH Para High (-4.81°) and RH Perp–LH Para High (-66.12°) conditions closely approximated the overall block slopes (-5.48° and -68.11°, respectively), indicating rapid adaptation. A similar pattern was observed at movement offset (Figure S3-B), where the group-level slopes in the second cycle in LH Perp–RH Para High (-7.70°) and RH Perp–LH Para High (-67.48°) conditions were reasonably close the overall group-level slopes (-8.47° and -69.88°, respectively). These findings suggest that rapid adjustments in movement direction in response to simultaneous gain perturbations emerged within the first 20 trials itself.

***Table S3:*** *Regression slopes for each 10-trial cycle and entire block of perpendicular deviations between the two arms in Experiment 3*

| | At Peak Velocity | | At Movement Offset | |
|---|---|---|---|---|
| **Cycle** | **LH Perp - RH Para High** | **RH Perp - LH Para High** | **LH Perp - RH Para High** | **RH Perp - LH Para High** |
| **1 (1 - 10 trials)** | -11.49° | -54.83° | -15.09° | -59.81° |
| **2 (11 - 20 trials)** | -4.81° | -66.12° | -7.70° | -67.48° |
| **3 (21 - 30 trials)** | -6.47° | -74.64° | -7.73° | -75.88° |
| **4 (31 - 40 trials)** | 0.04° | -76.00° | -5.34° | -74.64° |
| **5 (41 - 50 trials)** | -2.65° | -74.13° | -5.80° | -77.17° |
| **Entire Block** | -5.48° | -68.11° | -8.47° | -69.88° |

Table S4 shows the group-level slopes for each 10-trial cycle and the entire block for parallel deviations of the two arms at peak velocity and at movement offset. At peak velocity (Figure S4-A), the timescale of adaptation differed between the two asymmetric conditions. In the LH Perp–RH Para High condition, where the right arm had higher contribution to the parallel motion of the cursor, the slope in the third cycle (59.58°) surpassed the overall group-level slope (53.94°), and then oscillated around this value in subsequent cycles, suggesting a relatively fast adaptation followed by stabilization of the coordination strategy. In contrast, in the RH Perp–LH Para High condition, the slopes in the first four cycles fluctuated around the line of equivalence, showing no early shift. A small deviation was then observed in the fifth cycle (Slope = 39.78°), indicating a slower change in coordination when the left arm had higher parallel contribution. At movement offset (Figure S4-B), the group slope in both LH Perp–RH Para High (54.23°) and RH Perp–LH Para High (28.97°) conditions aligned closely with the respective overall group slopes (55.70° and 27.16°, respectively) by the second cycle, indicating rapid adaptation to the asymmetric gains.

***Table S4:*** *Regression slopes for each 10-trial cycle and entire block of perpendicular deviations between the two arms in Experiment 3*

| | **At Peak Velocity** | | **At Movement Offset** | |
|---|---|---|---|---|
| **Cycle** | **LH Perp - RH Perp High** | **RH Perp - LH Para High** | **LH Perp - RH Perp High** | **RH Perp - LH Para High** |
| **1 (1 - 10 trials)** | 48.80° | 46.37° | 48.31° | 29.54° |
| **2 (11 - 20 trials)** | 50.69° | 43.87° | 54.23° | 28.97° |
| **3 (21 - 30 trials)** | 59.58° | 42.41° | 57.69° | 29.36° |
| **4 (31 - 40 trials)** | 57.59° | 49.34° | 61.13° | 32.76° |
| **5 (41 - 50 trials)** | 53.33° | 39.78° | 63.28° | 18.55° |
| **Entire Block** | 53.94° | 44.13° | 55.70° | 27.16° |

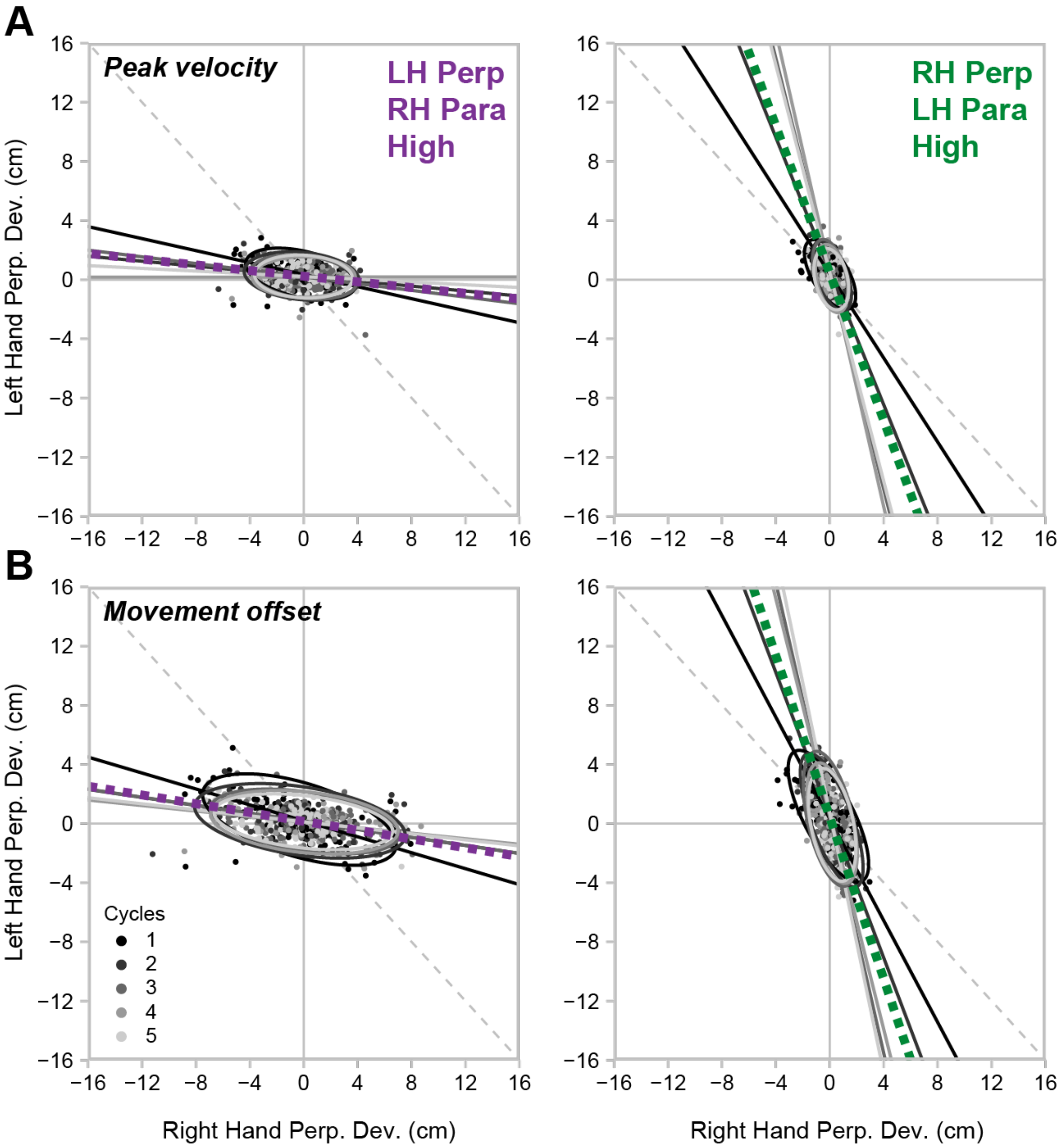


***Figure S3:*** *Group-level covariance relationships between the perpendicular deviations of the right and left arms over time in Experiment 3, shown separately for LH Perp–RH Para High (left column) and RH Perp–LH Para High (right column) conditions at **(A)** peak velocity and **(B)** movement offset. Each subplot shows orthogonal regression lines and 95% confidence ellipses for successive 10-trial cycles, with shading from dark to light grey indicating progression across the block. Coloured dotted lines represent the overall regression line fit for the entire block (50 trials) in each condition, identical to the group-level slopes in Figure 6D and 6E. Dashed grey line (slope = –45°) denotes the line of equivalence.*

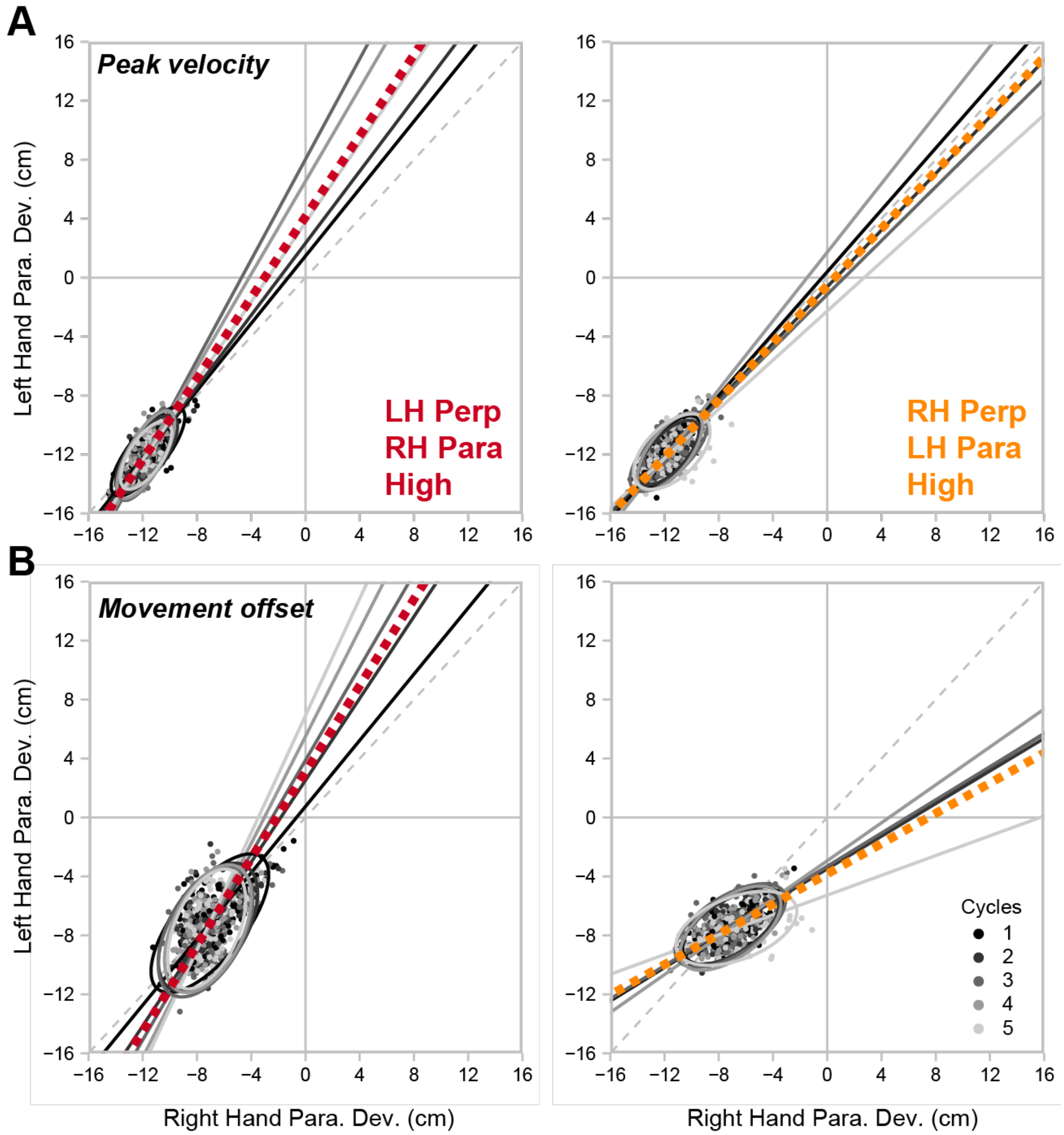


***Figure S4:*** *Group-level covariance relationships between the parallel deviations of the right and left arms over time in Experiment 3, shown separately for LH Para–RH Perp High (left column) and RH Para–LH Perp High (right column) conditions at **(A)** peak velocity and **(B)** movement offset. Each subplot shows orthogonal regression lines and 95% confidence ellipses for successive 10-trial cycles, with shading from dark to light grey indicating progression across the block. Coloured dotted lines represent the overall regression line fit for the entire block (50 trials) in each condition, identical to the group-level slopes in Figure 7D and 7E. Dashed grey line (slope = 45°) denotes the line of equivalence.*

**Experiment 4**

In experiment 4, we increased the contribution of one arm to the perpendicular and the parallel motion of the cursor, while decreasing the contribution of the other arm to both. To assess the timescale of changes in coordination across the perpendicular and parallel deviations of the two arms, we computed group-level covariance over successive 10-trial cycles separately for perpendicular and parallel deviations. Table S5 shows the group regression slopes for the perpendicular deviations of the two arms at peak velocity and at movement offset. At peak velocity (Figure S5-A), both LH Perp–Para High and RH Perp–Para High conditions showed early deviations from the Equal_1 slope, with slopes aligning with their respective overall group slopes by the second cycle (within 20 trials). This indicated quick adaptation evident during the early phases of the movement. In contrast, at movement offset (Figure S5-B), slope deviations from the Equal_1 condition were observed within the first 10 trials, but convergence to the overall block slopes occurred more gradually. By the third cycle, slopes in LH Perp–Para High (-22.86°) and RH Perp–Para High (-74.95°) closely matched their respective group-level slopes of the entire block (-21.61° and -74.42°), suggesting somewhat slower adaptation at movement end compared to earlier experiments.

***Table S5:*** *Regression slopes for each 10-trial cycle and entire block of perpendicular deviations between the two arms in Experiment 4*

| | At Peak Velocity | | At Movement Offset | |
|---|---|---|---|---|
| **Cycle** | **LH Perp - Para High** | **RH Perp - Para High** | **LH Perp - Para High** | **RH Perp - Para High** |
| **1 (1 - 10 trials)** | -35.99° | -69.00° | -29.38° | -67.93° |
| **2 (11 - 20 trials)** | -29.12° | -74.37° | -24.62° | -76.32° |
| **3 (21 - 30 trials)** | -24.49° | -70.92° | -22.86° | -74.95° |
| **4 (31 - 40 trials)** | -26.82° | -76.49° | -13.81° | -83.35° |
| **5 (41 - 50 trials)** | -22.40° | -73.68° | -11.79° | -74.18° |
| **Entire Block** | -27.92° | -72.59° | -21.61° | -74.42° |

Table S6 shows the group-level slopes for each 10-trial cycle as well as the entire block for the parallel deviations of the two arms at peak velocity and at movement offset. At peak velocity (Figure S6-A), no notable changes in slope from the Equal_1 condition were observed in the LH Perp–Para High condition for the first 40 trials, with only small deviations emerging by the fifth cycle (Slope = 36.64°). In contrast, in the RH Perp–Para High condition, consistent with the parallel deviation results from previous experiments, deviations in slopes were evident from the first cycle and progressively increased across cycles, surpassing the overall group-level slope by the fifth cycle (68.80°). This suggests more robust changes early on when the right arm made higher parallel contributions to cursor motion, while adaptation was slower and more gradual when the left arm had a higher contribution. However, at movement offset (Figure S6-B), both LH Perp–Para High and RH Perp–Para High conditions showed a clear deviation from the Equal_1 slope within the first 10 trials itself. Further, the slopes matched their respective overall group-level slopes by the third cycle (within 30 trials). By the fifth cycle, slope in the LH Perp–Para High condition decreased to 10.34° and the RH Perp–Para High slope increased to 85.93°. This suggested that participants continued to adjust the coordination strategies between the arms and increasingly constrained variability in the arm with higher contribution to the cursor, throughout the block.

***Table S6:*** *Regression slopes for each 10-trial cycle and entire block of perpendicular deviations between the two arms in Experiment 4*

| | At Peak Velocity | | At Movement Offset | |
|---|---|---|---|---|
| **Cycle** | **LH Perp - Para High** | **RH Perp - Para High** | **LH Perp – Para High** | **RH Perp - Para High** |
| **1 (1 - 10 trials)** | 44.42° | 58.38° | 20.85° | 73.76° |
| **2 (11 - 20 trials)** | 45.57° | 54.60° | 27.31° | 68.38° |
| **3 (21 - 30 trials)** | 43.49° | 58.97° | 17.71° | 74.36° |
| **4 (31 - 40 trials)** | 40.55° | 61.77° | 11.14° | 76.17° |
| **5 (41 - 50 trials)** | 36.64° | 68.80° | 10.34° | 85.93° |
| **Entire Block** | 41.90° | 60.12° | 16.32° | 75.50° |

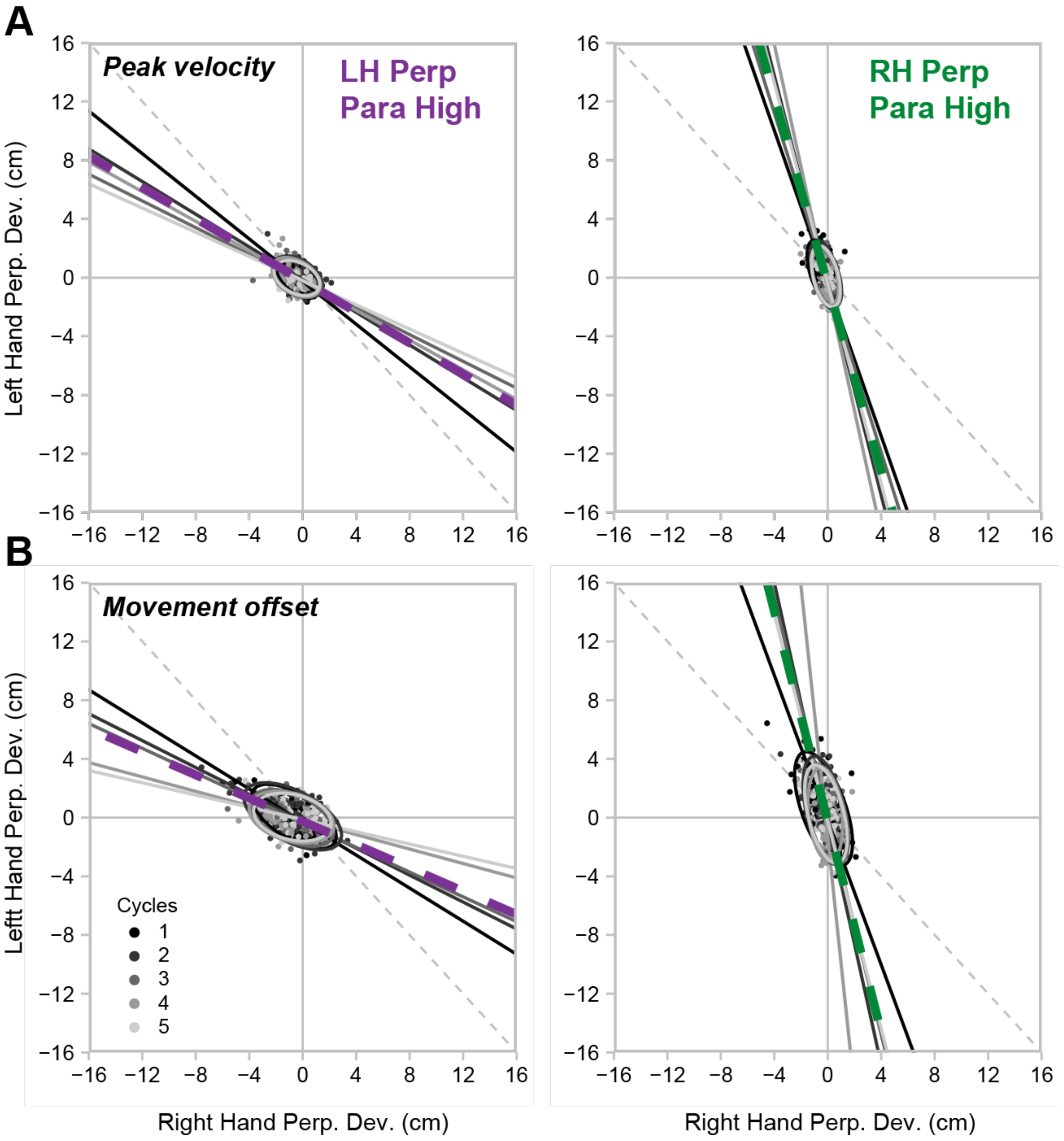


***Figure S5:*** *Group-level covariance relationships between the perpendicular deviations of the right and left arms over time in Experiment 4, shown separately for LH Perp–Para High (left column) and RH Perp–Para High (right column) conditions at* ***(A)*** *peak velocity and* ***(B)*** *movement offset. Each subplot shows orthogonal regression lines and 95% confidence ellipses for successive 10-trial cycles, with shading from dark to light grey indicating progression across the block. Coloured dotted lines represent the overall regression line fit for the entire block (50 trials) in each condition, identical to the group-level slopes in Figure 8D and 8E. Dashed grey line (slope = –45°) denotes the line of equivalence.*

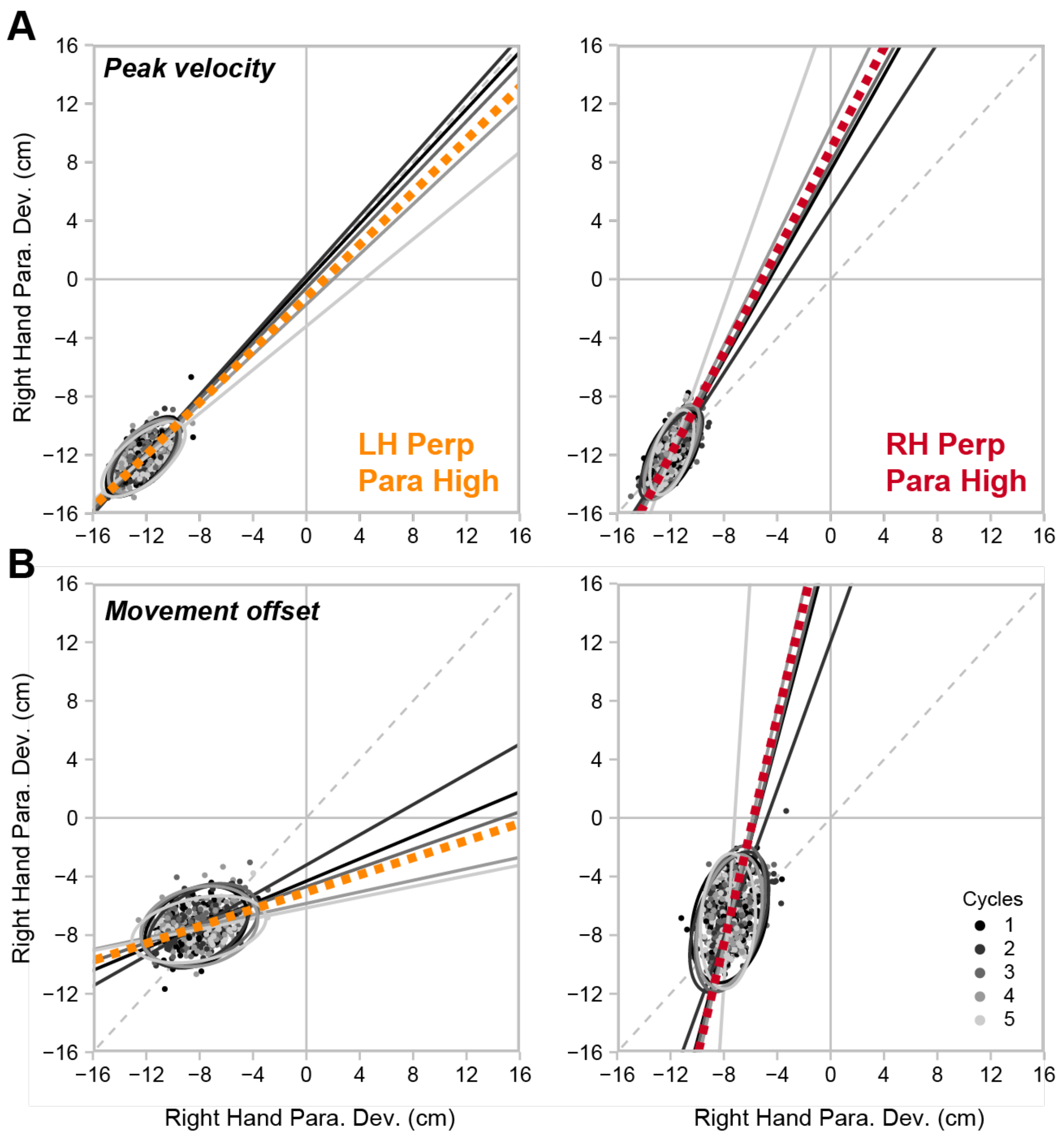


***Figure S6:*** *Group-level covariance relationships between the parallel deviations of the right and left arms over time in Experiment 4, shown separately for LH Perp–Para High (left column) and RH Perp–Para High (right column) conditions at **(A)** peak velocity and **(B)** movement offset. Each subplot shows orthogonal regression lines and 95% confidence ellipses for successive 10-trial cycles, with shading from dark to light grey indicating progression across the block. Coloured dotted lines represent the overall regression line fit for the entire block (50 trials) in each condition, identical to the group-level slopes in Figure 9D and 9E. Dashed grey line (slope = 45°) denotes the line of equivalence.*
